# Activation of the TP53 pathway is a therapeutic vulnerability in NUP98::KDM5A^+^ pediatric AML

**DOI:** 10.1101/2025.09.11.675580

**Authors:** Luca N. Cifarelli, Hasan Issa, Ludovica Proietti, Konstantin Schuschel, Lucie Gack, Katja Menge, Christian Ihling, Meike Vogler, Andrea Sinz, Jan-Henning Klusmann, Dirk Heckl

## Abstract

*NUP98::KDM5A*–rearranged pediatric acute myeloid leukemia (AML) is a rare, infancy-predominant entity with dismal outcome and no targeted therapeutic options. Given its suspected fetal origin, we hypothesized that leukemic maintenance depends on developmentally restricted vulnerabilities embedded within fetal hematopoietic programs. Using matched fetal and adult hematopoietic stem and progenitor cell models, we integrated transcriptomic and proteomic profiling with CRISPR–Cas9 screenings to define ontogeny-specific dependencies in NUP98::KDM5A leukemia. Fetal-derived NUP98::KDM5A leukemias exhibited greater *in vivo* aggressiveness, and retained fetal transcriptional signatures compared with adult counterparts. A comparative CRISPR–Cas9 screen using a library targeting fetal gene programs, conducted in both fetal- and adult-derived NUP98::KDM5A leukemias, identified the AAA⁺ ATPase TRIP13 as a selective and essential dependency in the fetal context. Mechanistically, TRIP13 physically interacted with the TP53 phosphatase PPM1D/WIP1, resulting in suppression of TP53 activation. Genetic ablation of *Trip13*, or pharmacologic inhibition using DCZ0415, a small-molecule inhibitor targeting TRIP13, reactivated TP53, induced G₂/M arrest and mitochondrial apoptosis, and depleted leukemic cells *in vitro*; these effects were fully rescued by *TP53* loss. In competitive transplantation assays, *Trip13* ablation significantly impaired leukemic fitness *in vivo*. Together, these findings define a developmentally restricted TRIP13–PPM1D–TP53 survival axis in NUP98::KDM5A AML and provide mechanistic proof-of-concept that reactivation of the TP53 signaling pathway via TRIP13 inhibition represents a therapeutically targetable vulnerability in this high-risk pediatric leukemia.

**Visual abstract:** 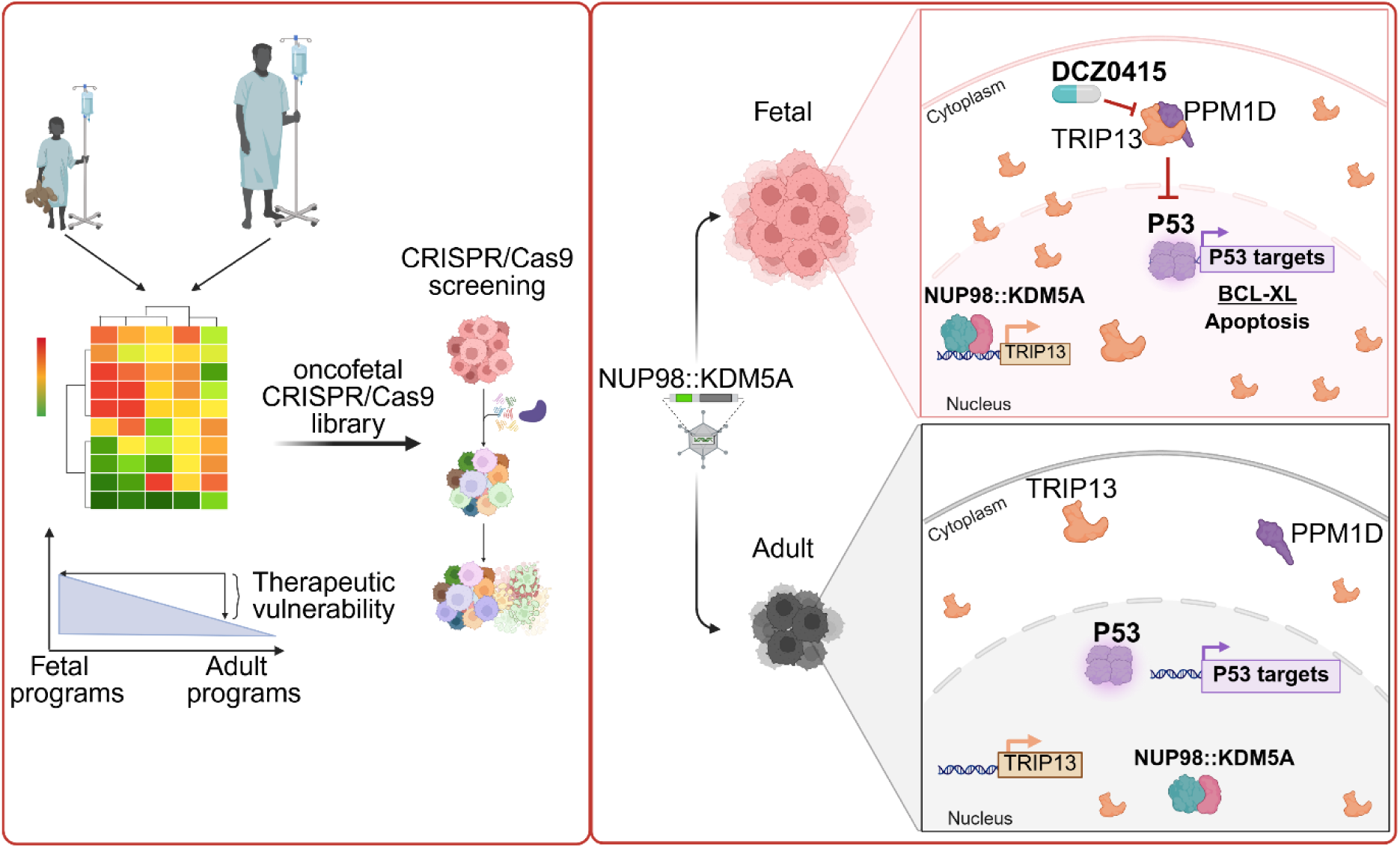

## Introduction

Pediatric acute myeloid leukemia (AML) is a biologically heterogenous disease, accounting for ∼15-20% of childhood acute leukemias, with a mutational landscape that differs substantially from adult AML and varies across pediatric age groups [1–5]. Increasing evidence suggests that this heterogeneity is shaped not only by genetic lesions but also by the developmental stage and cellular context in which transformation occurs, a concept that is particularly relevant in infant AML (≤2 years of age) [6–8]. In several pediatric AML subtypes, oncogenic driver lesions can be detected at birth, indicating an *in utero* origin and implicating fetal hematopoiesis as a permissive developmental context for leukemic initiation [9, 10]. Although direct causal links remain difficult to establish, the unique molecular properties of fetal hematopoietic cells and their niche are thought to contribute to the striking enrichment of specific chromosomal translocations and disease subgroups observed in early childhood.

Acute megakaryoblastic leukemia (AMKL) represents one such subgroup, characterized by aggressive clinical behavior and inferior event-free and overall survival in children without underlying Down syndrome (trisomy 21) [11–13]. Among pediatric AMKLs, the chromosomal translocation t(11;12) generates the NUP98::KDM5A fusion oncoprotein, composed of the N-terminal glycine-leucine-phenylalanine-glycine (GLFG) repeat motif of NUP98 and the C-terminal plant homeodomain finger (PHD3) domain of the histone lysine demethylase KDM5A (also known as JARID1A) [14, 15]. Expression of NUP98::KDM5A potently transforms murine hematopoietic stem/progenitor cells (HSPCs) and induces HOXA/B cluster gene activation, recapitulating core transcriptional and phenotypic features of the human disease [16–18].

While the fetal origin and developmental constraints of other high-risk pediatric AML entities, such as *CBFA2T3::GLIS2* and *KMT2A*-rearranged leukemias, have been extensively investigated [6], the contribution of fetal hematopoietic context to the initiation and maintenance of NUP98::KDM5A-driven leukemia remains largely unexplored [19–21]. Fetal liver HSPCs are highly proliferative and display a pronounced erythro-megakaryocytic bias, supporting rapid expansion of the lifelong hematopoietic stem cell pool through developmentally programmed epigenetic and transcriptional states, which renders them fundamentally distinct from their adult counterparts [22, 23]. These coordinated epigenetic landscapes and transcriptional programs raise the possibility that leukemias initiated in fetal hematopoiesis may remain dependent on developmentally restricted molecular circuits that are dispensable in adult cells.

Here we demonstrate that fetal hematopoietic origin markedly accelerates NUP98::KDM5A-driven leukemogenesis and enforces a critical TRIP13/PPM1D–TP53 survival axis. By exploiting this developmentally constrained dependency through genetic and pharmacologic inhibition of TRIP13, we uncover a mechanistically precise vulnerability that can be therapeutically engaged in this otherwise treatment-limited pediatric AML subtype.

## Materials and methods

### Cell culture

Human cell lines were purchased from the German Collection of Microorganisms and Cell Cultures (DSMZ) and maintained according to the supplier’s instructions. Fetal HSPCs were obtained from the MRC-Wellcome Trust Human Developmental Biology Resource (HDBR). Pediatric AML samples were collected from patients enrolled in the AML Berlin-Frankfurt-Münster treatment protocols for children and adolescents. The study was approved by the institutional review boards of all participating centers. All investigations were performed in accordance to the Declaration of Helsinki and local laws and regulations, and informed consent was obtained from all patients and custodians.

Human and murine HSPCs were prepared and cultured as previously described and maintained for at least 60 days [24, 25].

Media promoting the megakaryocyte differentiation of HSPCs consisted of respective media supplemented with corresponding cytokines: 15ng/mL SCF/Scf and 15ng/mL TPO/Tpo. All cytokines were purchased from Peprotech.

mFL-HSPCs were cultured in pre-stimulation medium composed by: DMEM supplemented with 10% FBS, 1% penicillin/streptomycin, 1% sodium pyruvate, 1% L-glutamine, 1% nonessential amino acids, 25ng/mL murine TPO and 25ng/mL murine SCF.

mBM-HSPCs were cultured in pre-stimulation medium composed by: IMDM supplemented with 10% FBS, 1% penicillin/streptomycin, 1% sodium pyruvate, 1% L-glutamine, 1% nonessential amino acids, 50ng/mL murine TPO and 50ng/mL murine SCF.

After 24 hours post-transduction, mFL-HSPCs and mBM-HSPCs were cultured in selection medium promoting megakaryocytic differentiation composed by: DMEM and IMDM, respectively, supplemented with 10% FBS, 1% penicillin/streptomycin, 1% sodium pyruvate, 1% L-glutamine, 1% nonessential amino acids, 15ng/mL murine Tpo and 15ng/mL murine Scf.

Human fetal and adult PB CD34+ HSPCs: cells were cultured in pre-stimulation medium composed by: Stemspan SFEM II supplemented with 1% penicillin/streptomycin, 0.75 µM Stemregenin, 35nM UM171, 100ng/ml human FLT3L, 50ng/ml human TPO, 100ng/ml human SCF, 20ng/ml human IL6. After 24 hours post-nucleoporation, hFL-HSPCs and adult PB CD34+ HSPCs cells were cultured in selection medium promoting megakaryocytic differentiation composed by: Stemspan SFEM II, respectively, supplemented with 10% FBS, 1% penicillin/streptomycin, 15ng/mL human TPO and 15ng/mL human SCF.

### Lentiviral vector construction and production

Lentiviral particles were generated by co-transfecting HEK293T cells with the corresponding expression constructs. pMD2.G and psPAX2 (Addgene #12259 and #12260), via the polyethyleneimine transfection method as previously described [26, 27]. Codon-optimized cDNAs were similarly cloned into the same SIN40C.SFFV backbone encoding GFP or dTomato as a fluorescent reporter [24, 25]. For inducible expression, a doxycycline-inducible SIN40C.TRE vector was used [25]. All plasmids generated in this study have been deposited at Addgene and are listed in Supplementary Table 1.

A list of sgRNAs is in Supplementary Table 2 (designed using CCTop online tools [28]) that have been cloned into the SGL40C.EFS.E2Crimson or SGL40C.EFS.dTomato vectors (Addgene #100894 and #89395) [29].

NUP98::KDM5A encoding cDNA was designed based on a report in human patients [30]. TRIP13-encoding cDNA was designed based on Ensembl genome browser. All cDNAs were tagged at the 5’ terminus with three HemoAgglutinin tag (3xHA), flanked with AgeI at 5’ terminus and MluI at 3’ terminus, codon optimized and ordered from pTwistAmp.

Plasmids were retransformed and cDNAs were subcloned using the restriction enzymes AgeI and MluI into the backbone vectors for constitutive expression: NUP98::KDM5A cDNA in SIN40C.SFFV.GFP.IRES.MCS and TRIP13 in cDNA in SIN40C.SFFV.MCS.IRES.dtomato and in SIN40C.TRE.MCS.IRES.dTomato.PGK.sfGFP.P2A.Tet3G.

### Transduction

Cells were transduced with concentrated viral particles in the presence of 2-5µg/ml polybrene as previously described [24, 25, 31]. Primary human and murine HSPCs were transduced in Retronectin-coated plates according to manufacturer’s instructions (Takara Bio) as previously described [31].

### CRISPR/Cas9 library screening

15 x 10^6^ fetal and adult NUP98::KDM5A^+^, FPD-AML and ML-DS models were lentivirally transduced with the fetal signature sgRNA library (MOI=0.3) to achieve sufficient representation (1000-fold coverage per sgRNAs). Samples were harvested at 2 days post-transduction and at 16 population doublings, corresponding to approximately 30 days of culture. Genomic DNA was isolated as previously described [24].

704 genes of our CRISPR-Cas9 library, targeting the TP53 pathway, were collected using publicly available GSEA gene sets.

Model-based analysis of genome-wide CRISPR/Cas9 knockout (MAGeCK) [32] was used to identify hits from the sgRNA screen. Double barcoded reads were demultiplexed using custom R scripts; guides with fewer than 20 reads in ≥75% of all samples were excluded. Raw read counts were passed to the MAGeCK test command using default parameters. Samples collected at the endpoint were compared to day 2 to determine negative enrichment. Genes with a P-value <0.05 as determined by MAGeCK were deemed significant.

### Proliferation and rescue assays

To evaluate the effect of NUP98::KDM5A transformation, mFL-HSPCs and mBM-HSPCs were co-transduced with lentiviral vectors overexpressing the human cDNA encoding NUP98::KDM5A, which was assigned to a green fluorescent protein thereby allowing direct clonal growth competition.

To assess growth advantages conferred by overexpression of sgRNA(s) or TRIP13 cDNA and for rescue experiments, transduced mFL-HSPCs were measured every 2/3 days starting 72 hours post-transduction and analyzed for reporter gene expression. For inducible expression of TRIP13, doxycycline was added to culture medium at a concentration of 0.5 μg/mL 96 hours post transduction.

### Flow cytometry and sorting

To assess the expression of fluorescent reporter protein and their immunophenotype, cells were analyzed for the expression of surface cell markers using fluorochrome-coupled antibodies, listed in Supplementary Table 1. The following panels were used: Myeloid (CD11b and Gr-1), Progenitor (CD34, CD16/32, C-kit/CD117, Sca-1), Megakaryocytic (CD41, CD42d) Erythroid (CD71 and CD235a for human cells or Ter119 for murine cells). Flow cytometry was performed on a CytoFLEX flow cytometer (Beckman Coulter) and the data analysis was done using Kaluza 1.5 (Beckman Coulter) software.

The same procedure was performed to sort the cell subpopulation of interest. Cells were sorted using BD FACSAria™ II Flow Cytometer (BD Biosciences).

Apoptotic cell death and cell cycle analysis was performed as previously described [25].

### Micrographs

Micrographs were obtained with a Keyence BZ-9000 Microscope using the BZ-II viewer and images were processed with the BZ-II Analyzer software.

### Drug response assays

For drug response curves using DCZ0415 (MedChemExpress), cell viability was assessed using the CellTiter-Glo® Cell Viability Assay (Promega) according to the manufacturer’s instructions after 2-5-7 days post-treatment.

Synergy plots, summary Bliss score calculations were generated and performed with SynergyFinder [33]. All conditions were normalized to DMSO treated control. Data represent the mean ± SD of biological triplicates.

### Proteomic analysis

Total cell lysis and Western blotting were performed as previously described [24, 25]. Blots were developed using Amersham™ ECL Prime Western Blotting Detection Reagent (Thermo Fisher Scientific).

Co-immunoprecipitation of TRIP13 was performed on OCI-AML3 cells using Anti-HA antibody coupled to magnetic beads (Pierce Anti-HA Magnetic Beads). After cell lysis and affinity pulldown, proteins were processed as previously described [24, 25].

### Gene expression profiling

NUP98::KDM5A and control (EV) transduced mFL-HSPCs and mBM-HSPCs were FACS-sorted 3, 21 days after transduction; sgLuc- and sgTrip13- transduced mFL-HSPCs expressing NUP98::KDM5A translocation were FACS-sorted 4 days after transduction. Fetal NUP98::KDM5A cells were treated with DCZ0415 or DMSO control 24h prior gene expression profiling. RNA was extracted using the Quick-RNA Microprep Kit (Zymo Research) as previously described [25]. A minimum amount of 150ng RNA was used as input material for the RNA sample preparations. RNA-sequencing was performed by Novogene Company, Ltd. Sequencing libraries were generated and sequenced as previously described [25].

Raw FASTQ data (raw reads) were first pre-processed using fastp [34] and then further processed as previously described [35]. Differential expression analysis was performed using the DESeq2 package in R [36]. The resulting P-values were adjusted using the Benjamini and Hochberg’s approach for controlling the False Discovery Rate (FDR) [37]. Genes with an adjusted P-value <0.05 were considered differentially expressed.

Gene set enrichment analysis (GSEA) was performed using the Broad GSEA software [38, 39] with the permutation type set to “Gene_set” (1000 permutations). Human gene symbols were mapped to murine gene symbols using orthologue annotations provided by Ensembl [40] considering only one-to-one orthologue relationships.

### Animal studies

All experiments involving mice were performed in accordance with protocols approved by the local authorities (Landesverwaltungsamt Sachsen-Anhalt).

B6J.129 (Cg)-Gt(ROSA)26Sortm1.1(CAG-cas9*.-EGFP)Fezh/J) (Jackson Laboratory. RRID: IMSR_JAX:026179) and C57BL/6J (Charles River. RRID: IMSR_JAX:000664) mice were maintained in a specific pathogen-free environment in individual ventilated cages and fed with autoclaved food and water at the Martin-Luther-University Halle-Wittenberg.

### Transplantations and analysis of leukemic mice

2 x 10^6^ cells were injected intravenously into sublethally irradiated (7 Gy) C57BL/6N mice. Engraftment of the transplanted cells was evaluated by tail bleeding of the recipients every 4 weeks. When signs of disease occurred, the mice were sacrificed and harvested bone marrow cells were stained with the following fluorochrome-coupled antibodies describe above.

### Statistical analysis

All statistical analyses were performed with GraphPad Prism 8 (San Diego, CA). Unless indicated otherwise, 2-tailed Student’s t-tests were used to determine the P-value between 2 groups. Statistical analyses of experiments were performed with analysis of variance (ANOVA) with Bonferroni *post hoc* analysis for comparisons between ≥3 groups. Kaplan-Meier method was used to determine mice survival using log-rank (Mantel-Cox) tests for statistical analyses. *P, .05, **P, .01, ***P, .001, ****P, .0001. A value of P, .05 was considered statistically significant.

All statistical tests and sample numbers are disclosed in the respective figure legends and supplemental tables.

## Results

### NUP98::KDM5A induces highly aggressive leukemia in fetal HSPCs

To investigate whether the developmental origin of the leukemia initiating cell is an accelerator of NUP98::KDM5A-driven leukemogenesis, we established matched fetal and adult murine leukemia models. We transduced murine hematopoietic stem and progenitor cells (HSPCs) from fetal liver at embryonic day 13.5 (mFL-HSPCs) and from adult bone marrow (mBM-HSPCs) with a lentiviral vector expressing human NUP98::KDM5A cDNA [30] (Fig. 1a). Both fetal- and adult-derived HSPCs exhibited progressive expansion of NUP98::KDM5A^+^ cells, whereas control vector–transduced cells exhausted over time, indicating efficient leukemic transformation in both developmental contexts (Fig. 1b and Supplementary Fig. 1a). However, despite comparable transduction efficiencies and similar baseline immunophenotype, fetal-derived NUP98::KDM5A^+^ cells expanded significantly faster than their adult counterparts, consistent with an ontogeny-dependent increase in leukemic fitness (Fig. 1b and Supplementary Fig. 1a).

**Fig. 1.**
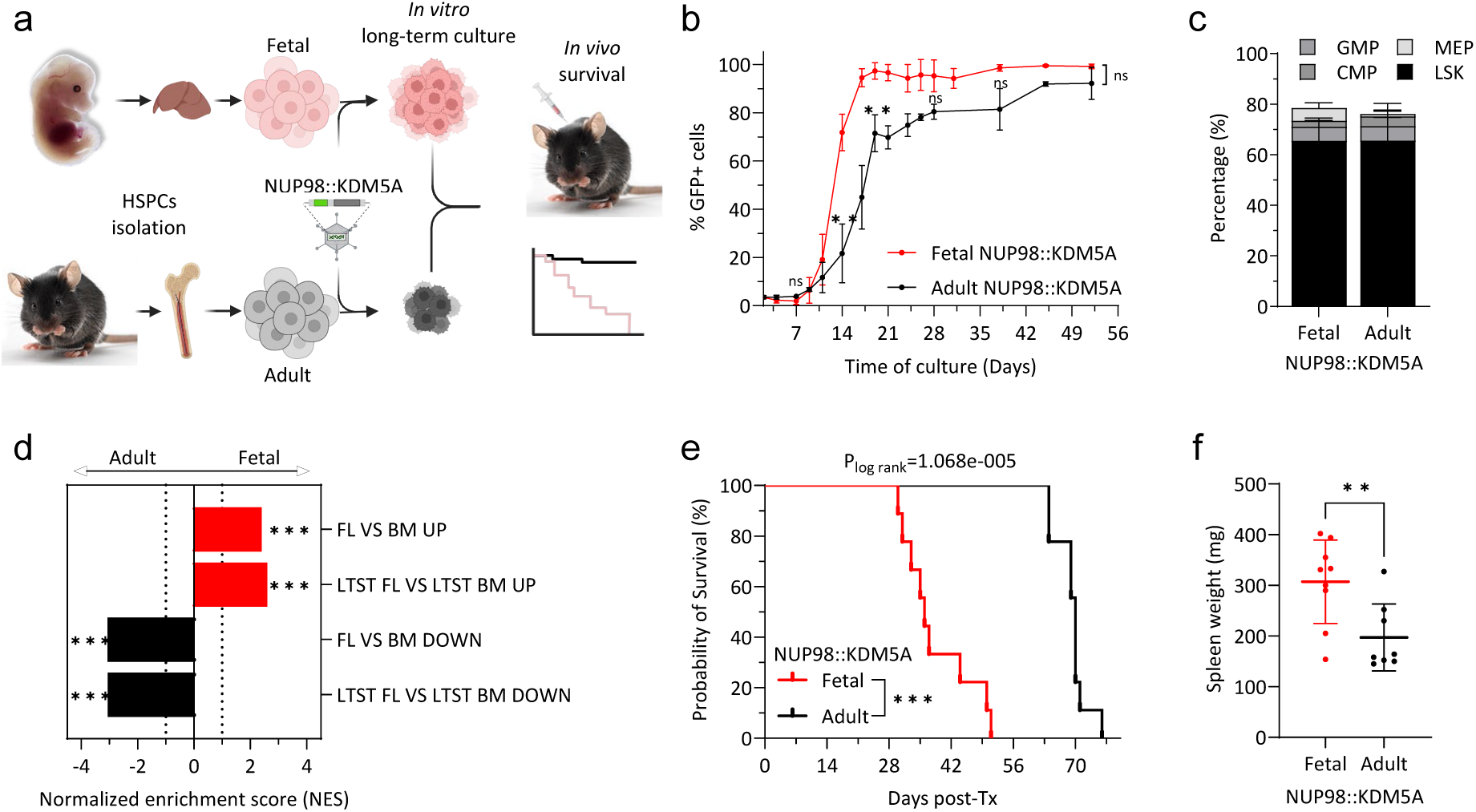
NUP98::KDM5A fusion oncoprotein induces a more aggressive disease in fetal cells. a) Schematic workflow for the generation of in vitro and in vivo murine models of constitutive overexpression of NUP98::KDM5A fusion oncoprotein in murine Hematopoietic Stem and Progenitor Cells (HSPCs) isolated from fetal liver (mFL-HSPCs) of healthy fetuses at E13.5 and bone marrow of adult mice (mBM-HSPCs). Created in Biorender.com b) GFP+ percentage of fetal and adult NUP98::KDM5A+ cells maintained in liquid culture. Data from 1 representative experiment performed in replicates (n > 3) are shown as mean s.d. (t-test). c) Bar graph showing the percentage of LSK (Lineage negative, c-kit+ and Sca1+), Megakaryocytic-erythroid (MEP; Lineage negative, c-kit+, Sca1-, CD16/CD32- and CD34+) and common myeloid (CMP; Lineage negative, c-kit+, Sca1-, CD16/CD32- and CD34-) and granulocyte/monocyte (CMP; Lineage negative, c-kit+, Sca1-, CD16/CD32+ and CD34+) progenitors of fetal and adult NUP98::KDM5A+ cells after 4 weeks of culture. (mean ± s.d., n > 3) d) Bar graphs showing the normalized enrichment scores (NES) of significantly upregulated or downregulated gene sets in fetal NUP98::KDM5A^+^ cells compared to adult NUP98::KDM5A^+^ cells. e) Kaplan-Meier survival curves of C57BL/6N mice transplanted with fetal and adult NUP98::KDM5A^+^ cells (n ≥ 6). f) Dot plot showing spleen weights of C57BL/6N mice transplanted with fetal and adult NUP98::KDM5A^+^ cells (mean ± s.d., n ≥ 6) (t-test). ns, not significant * P < .05, ** P < .01, ***P < .001, ****P < .0001

Immunophenotypic analysis showed that NUP98::KDM5A expression maintained cells in a highly immature LSK (Lineage⁻ c-Kit⁺ Sca1⁺) state with marked maturation arrest, findings that were corroborated by transcriptional profiling (Fig. 1c, Supplementary Fig. 1b-c and Supplementary Table 3). Gene expression analysis further revealed that fetal NUP98::KDM5A^+^ cells retained transcriptional signatures associated with fetal hematopoietic origin compared to adult-derived cells, highlighting the influence of developmental origin on leukemic phenotype and aggressiveness (Fig. 1d and Supplementary Table 4). Consistent with patient samples, both models exhibited upregulation of *HOXA* and *HOXB* gene clusters, together with marked downregulation of *RB1* target genes (Supplementary Fig. 1c and Supplementary Table 3), resembling a transcriptional program of the RB1-mutant state frequently associated with NUP98::KDM5A AML [12, 15, 41].

*In vivo*, transplantation of fetal NUP98::KDM5A^+^ cells into syngeneic recipients resulted in significantly accelerated leukemia onset compared to adult-derived cells (median disease-free survival: fetal 36 days versus adult 70 days; log-rank *P* = 1.07 × 10⁻⁵; Fig. 1e). Splenomegaly was significantly more pronounced in mice transplanted with fetal NUP98::KDM5A^+^ cells (Fig. 1f). Leukemic blasts in the bone marrow expressed high levels of the mature myeloid surface marker Gr-1 and CD11b (Mac-1), together with persistence of an immature LSK population (Supplementary Fig. 1d-e).

Collectively, these data demonstrate that developmental origin is a major determinant of NUP98::KDM5A-driven leukemic aggressiveness, with fetal hematopoietic cells providing a permissive context that accelerates disease onset and progression.

### Fetal dependency screening reveals TRIP13 as a critical vulnerability in NUP98::KDM5A^+^ AML

The aggressive phenotype of fetal-derived NUP98::KDM5A leukemia is closely linked to the persistence of fetal transcriptional programs, which remained active in a post-fetal hematopoietic environment. We reasoned that such developmentally regulated programs might impose context-specific dependencies required for leukemic maintenance and therefore represent a potential therapeutic vulnerability. To identify these dependencies, we performed a CRISPR-Cas9 loss-of-function screen targeting 867 genes, overexpressed in both human and murine fetal liver HSPCs compared to their adult counterparts [42], in fetal- and adult-derived NUP98::KDM5A^+^ leukemia models (Fig. 2a-b). To distinguish NUP98::KDM5A- and fetal-specific dependencies from broader AML requirements, we complemented these screens with two additional murine AML models: familial-platelet disorder with predisposition to AML (FPD-AML, loss of *Runx1* and *Bcor*) [43], and myeloid leukemia in Down syndrome (ML-DS) (Fig. 2c and Supplementary Fig. 2a) [24, 31, 44]. Applying a depletion threshold of >70% identified seven candidates, with the the AAA+ ATPase (ATPases Associated with diverse cellular Activities) *Trip13* displaying the highest fetal-to-adult dependency ratio (Supplementary Fig. 2b). The effect of *Trip13* knock-out was subsequently validated in a single-well fluorescence-based depletion assay using independent sgRNAs, confirming robust on-target effects (Fig. 2d). To corroborate these findings in a human system, we performed CRISPR-based *TRIP13* knock-out in a panel of human leukemic and non-leukemic (HT1080) cell lines, and primary human fetal HSPCs expressing NUP98::KDM5A. Consistent with the murine results, *TRIP13* loss markedly impaired the growth of NUP98::KDM5A⁺ human fetal HSPCs, whereas most control cell lines showed only modest proliferation defects (Fig. 2e). Gene expression analyses further demonstrated that *TRIP13* is upregulated as part of a fetal transcriptional program, with higher expression in fetal compared to adult HSPCs, and in pediatric AMKL relative to adult HSPCs and mature hematopoietic populations (Fig. 2f). Notably, the NUP98::KDM5A fusion oncoprotein not only maintained but further increased *TRIP13* expression in fetal but not in adult cells, indicating a fetal context-specific effect of the fusion on *TRIP13* regulation (Fig. 2g).

**Fig. 2.**
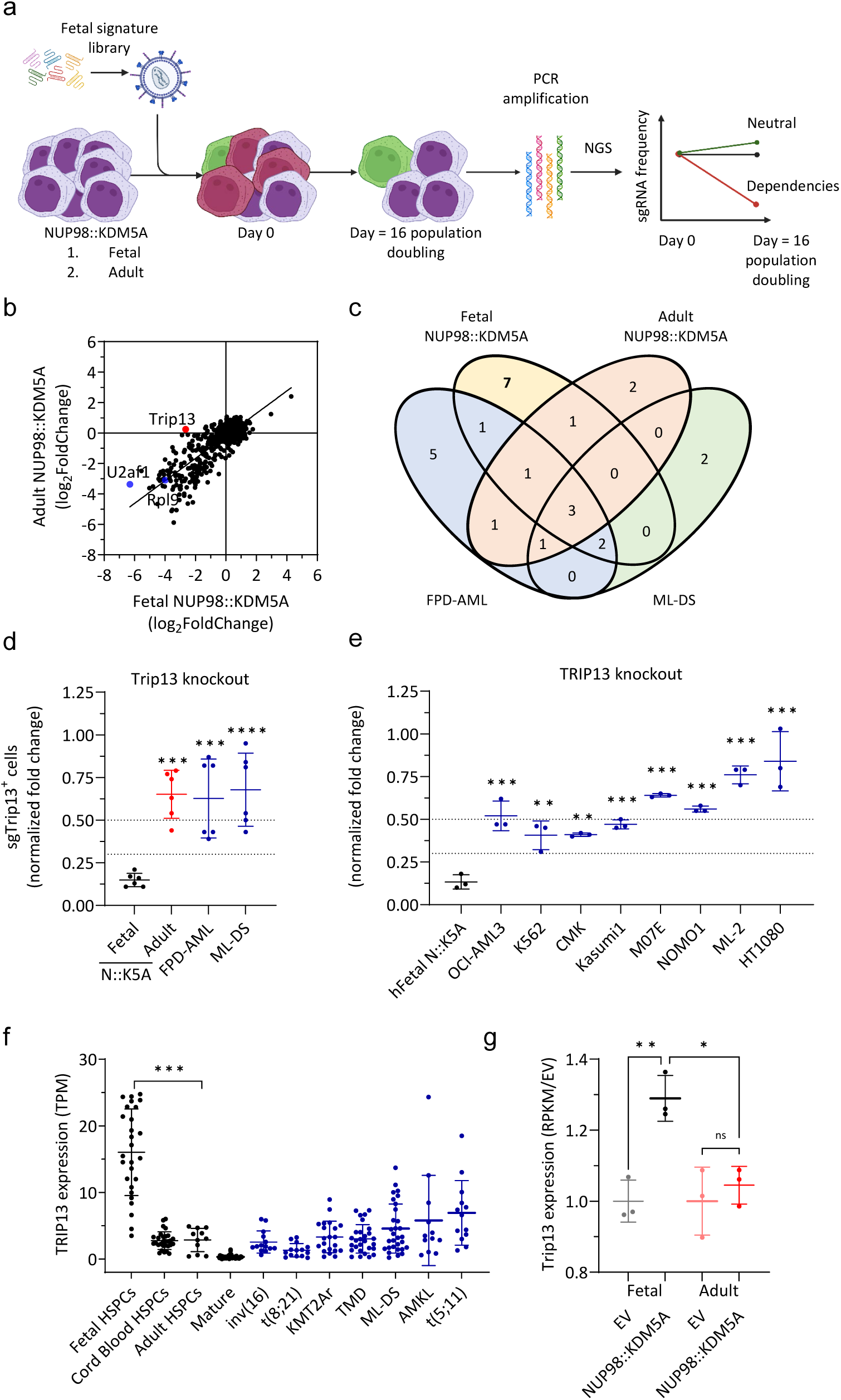
CRISPR-Cas9 knock-out screening reveals *TRIP13* dependency in fetal NUP98::KDM5A^+^ model. a) Schematic workflow of the high-throughput fetal signature-based CRISPR-Cas9 *in vitro* screen. b) Dot plots showing the log2 fold change of the fetal enriched genes targeted by sgRNAs in fetal (x-axis) and adult (y-axis) NUP98::KDM5A^+^ models. Blue dots represent positive control (*Rpl9*, *U2af1*). Red dots represent selected candidate gene *Trip13*. (n =2) c) Venn diagram illustrating significantly (P-value < 0.05) depleted (depletion > 70%) sgRNAs in different murine leukemia models. d) Dot plot showing the percentage of *Trip13*-targeting sgRNA-transduced fetal, adult NUP98::KDM5A^+^ (N::K5A), FPD-AML and ML-DS models after 21 days of culture, normalized to (negative control sgRNA [sgCtr]) and to day 2 ( mean ± s.d., n > 3 per sgRNA, 1-way ANOVA). e) Dot plot showing the percentage of *TRIP13*-targeting sgRNA-transduced hFL NUP98::KDM5A^+^ (N::K5A) cells and leukemia cell lines (OCI-AML3; K562; CMK; Kasumi1; M07E; NOMO1; ML-2) and non-leukemia cell line (HT1080) after 21 days of culture, normalized to (negative control sgRNA [sgCtr] and positive control sgRNA [sgRPL9 and sgU2AF1]) and to day 2 (mean ± s.d., n = 3 per sgRNA, 1-way ANOVA). f) Dot plot showing the expression of *TRIP13* in subgroups of HSPCs and pediatric AML from the HemAtlas dataset. g) Dot plot showing the expression of *Trip13* in control (EV) or NUP98::KDM5A^+^ murine fetal and adult cells after 21 days of culture (n = 3). ns, not significant * P < .05, ** P < .01, ***P < .001, ****P < .0001.

Collectively, these data identify *TRIP13* as a fetal context-dependent vulnerability in NUP98::KDM5A^+^ leukemias, supporting its prioritization for mechanistic and therapeutic interrogation in this high-risk AML subtype.

### Trip13 loss triggers TP53-dependent apoptosis

To further confirm that *Trip13* loss was causative for the observed growth defect, we co-transduced fetal NUP98::KDM5A⁺ cells with *TRIP13* cDNA together with *Trip13*-targeting sgRNAs. *TRIP13* overexpression fully rescued cell depletion, demonstrating on-target specificity of *Trip13* disruption (Fig. 3a). *Trip13* knock-out was confirmed by Western blotting (Supplementary Fig. 3a), and resulted in a marked increase in apoptosis (Fig. 3b) and a G₂/M cell-cycle arrest (Fig. 3c), consistent with the established role of TRIP13 in mitotic checkpoint control [45]. *Trip13* loss did not alter the differentiation status of leukemic cells, as assessed by immunophenotypic analysis (Supplementary Fig. 3b).

**Fig. 3.**
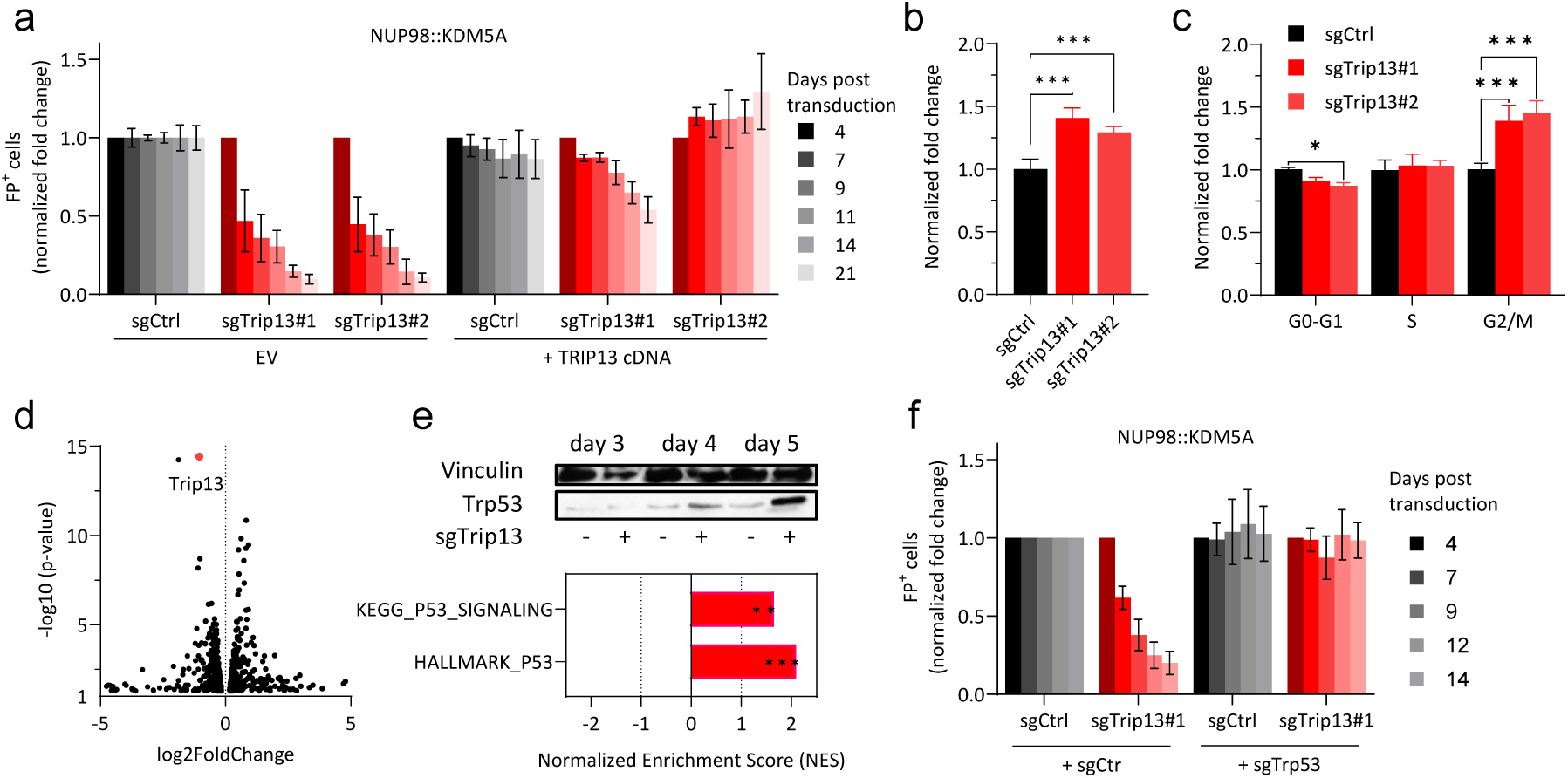
*Trip13* loss activates the TP53 pathway and arrests fetal NUP98::KDM5A^+^ cell proliferation. a) Bar plot showing the percentage of sgRNA-transduced (negative control sgRNA [sgCtr], sgTrip13#1-2) empty vector (EV) and TRIP13 overexpressing fetal NUP98::KDM5A^+^ cells after 14 days of culture, normalized to day 4 (mean±s.d., n > 3 per sgRNA, 1-way ANOVA) b) Bar graphs showing the percentage of Annexin-V^+^ sgRNA-transduced (negative control sgRNA [sgCtr], sgTrip13#1-2) fetal NUP98::KDM5A^+^ cells after 5 days of culture, normalized to control sgRNA (sgCtr) (mean ± s.d., n > 3 per sgRNA, 1-way ANOVA). c) Bar plot showing the percentage of sgRNA-transduced (negative control sgRNA [sgCtr], sgTrip13#1-2) fetal NUP98::KDM5A^+^ cells in G0-G1, S and G2/M phase after 5 days of culture (mean ± s.d., n > 3 per sgRNA, 2-way ANOVA). d) Volcano plot showing the differentially expressed genes upon *Trip13* knock-out in fetal NUP98::KDM5A^+^ cells P-value < 0.05 (n = 3). RNA-seq-based gene expression analysis of sgRNA-transduced (negative control sgRNA [sgCtr], sgTrip13#1) fetal NUP98::KDM5A^+^ cells was performed after 4 days of culture. e) Timeline Western blot showing TP53 protein levels in sorted sgRNA-transduced (negative control sgRNA [Luc], sgTrip13#1) fetal NUP98::KDM5A^+^ cells 4 days after transduction. Endogenous Vinculin was used as loading control. Bar graphs showing the normalized enrichment scores (NES) of significantly deregulated gene sets associated with TP53 pathway in *Trip13* knock-out fetal NUP98::KDM5A^+^ cells compared to control sgRNA. f) Bar plot showing the percentage of sgRNA-transduced (negative control sgRNA [sgCtr], sgTrip13, sgTrp53) fetal NUP98::KDM5A^+^ cells after 14 days of culture, normalized to day 4 ( mean±s.d., n > 3 per sgRNA, 1-way ANOVA) ns, not significant * P < .05, ** P < .01, ***P < .001, ****P < .0001.

To define the mechanism underlying *Trip13* dependency, we profiled transcriptomic changes four days after CRISPR-Cas9-mediated *Trip13* deletion in fetal NUP98::KDM5A^+^ cells (Fig. 3d). Gene set enrichment analysis (GSEA [38]) revealed a striking and selective enrichment of TP53 signaling pathway signatures upon *Trip13* loss (Fig. 3e and Supplementary Fig. 3c, Supplementary Table 5). In line with this transcriptional response, TP53 protein levels increased in a time-dependent manner after *Trip13* knock-out (Fig. 3e). Importantly, concomitant deletion of *Trp53* rescued the growth defect induced by *Trip13* loss (Fig. 3f), demonstrating that TP53 activation is required for the *Trip13*-dependent loss of leukemic cell fitness in fetal NUP98::KDM5A^+^ AML cells. Given the reported role of TRIP13 in DNA damage repair [46, 47], we next examined whether TP53 activation resulted from increased genotoxic stress. Although leukemic cells exhibited baseline γH2AX positivity, *Trip13* deletion did not further augment γH2AX levels compared with controls (Supplementary Fig. 3d), arguing against DNA damage-accumulation as the primary trigger of TP53 activation in this context.

Together, these results establish that fetal *NUP98::KDM5A⁺* AML cells are sensitive to TP53 activation and that *Trip13* sustains leukemic survival by suppressing TP53 activity, without requiring increased DNA damage signaling.

### TRIP13 interacts with PPM1D/WIP1 to repress TP53 pathway activation

To elucidate how TRIP13 regulates TP53 activity, we next mapped TRIP13 protein-protein interaction network using co-immunoprecipitation followed by mass spectrometry (IP-MS). Doxycycline-inducible hemagglutinin (HA)-tagged TRIP3 was expressed in a TP53 wild-type leukemia cell line (OCI-AML3) (Fig. 4a) [48, 49]. Among the proteins identified by TRIP13 pull-down, we focused on 33 candidates previously annotated in the STRING interaction database to be functionally or physically connected to TP53 (Fig. 4b). To prioritize TRIP13-associated complexes and identify the key mediators of TP53 suppression, we reasoned that the loss of a critical TRIP13 interactor would phenocopy *Trip13* deletion. Specifically, we predicted that disruption of such an interactor would induce TP53-dependent depletion of fetal NUP98::KDM5A⁺ cells, a phenotype reversible by concomitant *Trp53* deletion. Consistent with this hypothesis, CRISPR-mediated knock-out of 16 of the 33 candidate interactors resulted in a depletion pattern similar to that observed following *Trip13* knock-out (Supplementary Fig. 4a), which could be rescued by *Trp53* knock-out in 6 of the 16 TRIP13-associated candidates (Supplementary Fig. 4b). Among those 6 interactors, the protein phosphatase PPM1D/WIP1, a well-known regulator of TP53, emerged as the top candidate fulfilling both criteria: induction of leukemic cell depletion upon loss and rescue of this phenotype by *Trp53* co-deletion (Fig. 4c and Supplementary Fig. 4b). PPM1D/WIP1 is a well-established regulator of TP53 signaling, acting both directly and indirectly, including stabilization of MDM2 [50]. Co-immunoprecipitation followed by Western blotting confirmed a physical interaction between TRIP13 and PPM1D/WIP1 (Fig. 4d), and consistent with a functional interaction, deletion of either *Trip13* and *Ppm1d* in fetal NUP98::KDM5A^+^ cells resulted in increased TP53 protein levels (Fig. 4e). Notably, *PPM1D/WIP1* expression, similar to *TRIP13*, was elevated in pediatric AMKL patients compared to adult HSPCs and mature hematopoietic cells (Supplementary Fig. 4c), supporting the relevance of this regulatory axis in pediatric disease.

**Fig. 4.**
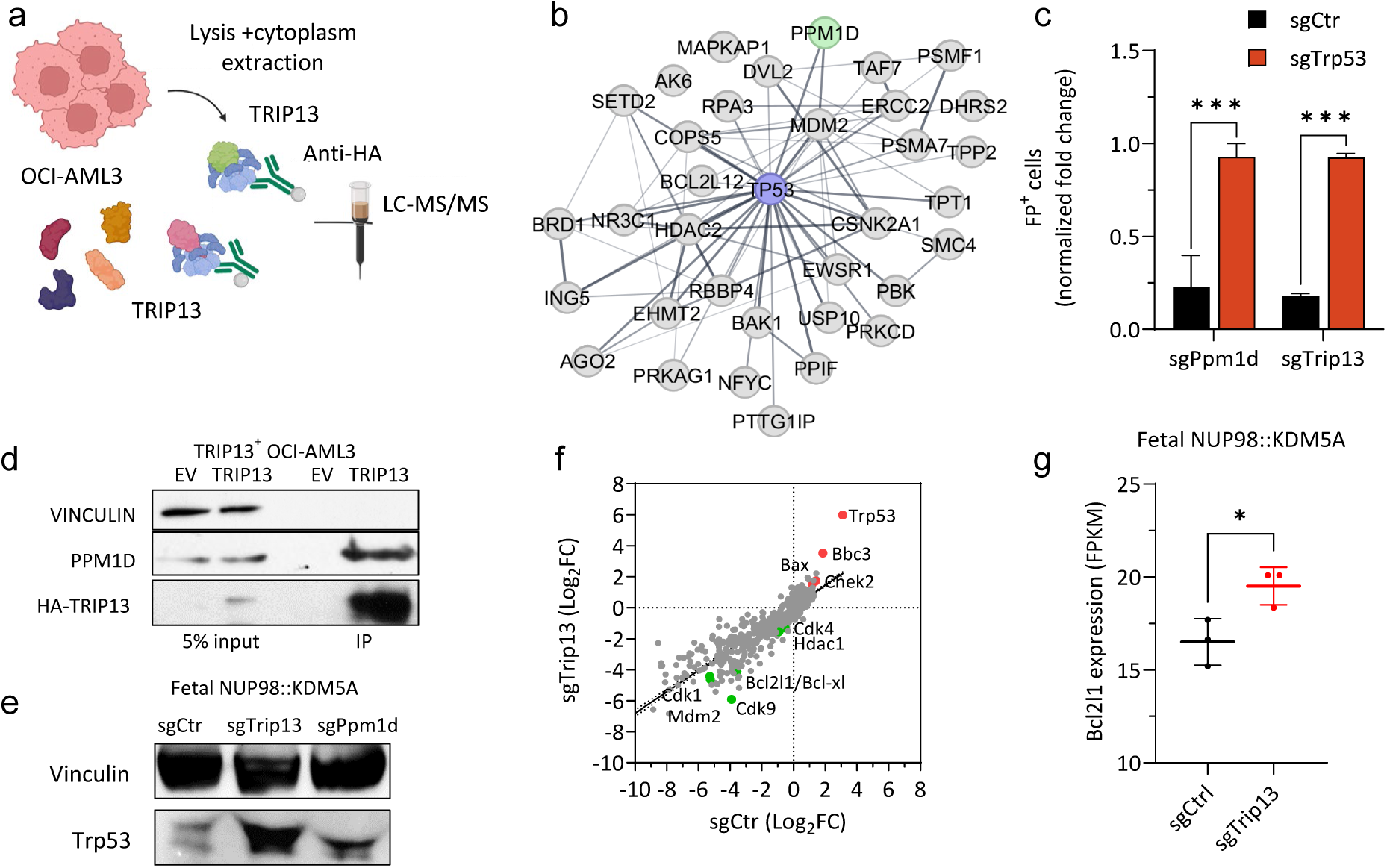
TRIP13 interacts with PPM1D/WIP1 to repress the TP53 pathway. a) Experimental workflow for isolating HA-tagged TRIP13 protein and its interactors from OCI-AML3 cells. b) String protein-protein interaction network of 33 TRIP13-interacting candidates. Network nodes (colored circles) represent proteins, lines between the nodes (edges) indicate the different types of interaction (functional and physical protein associations), the line thickness indicates the confidence of interaction (0.4). c) Bar plot showing the percentage of sgRNA-transduced (sgPpm1d, sgTrip13) fetal NUP98::KDM5A^+^ cells in combination with negative control sgRNA [sgCtr] or sgTrp53 after 14 days of culture, normalized to day 2 ( mean±s.d., n > 3 per sgRNA, 1-way ANOVA) . d) Western blot showing co-immunoprecipitation of PPM1D after HA-pulldown of HA-tagged TRIP13 in OCI-AML3 cells. Endogenous Vinculin was used as loading control. e) Western blot showing TP53 protein levels in sorted sgRNA-transduced (negative control sgRNA [Luc], sgTrip13 and sgPpm1d) fetal NUP98::KDM5A^+^ cells 4 days after transduction. Endogenous Vinculin was used as loading control. f) Dot plots showing the log2 fold change of the TP53 pathway-related genes targeted by control (x-axis), *Trip13*-depleted (y-axis) fetal NUP98::KDM5A^+^ cells. Red and green dots represent selected candidate genes. (n = 2) g) Dot plot showing the expression of *Bcl2l1* of control or *Trip13*-depleted fetal NUP98::KDM5A^+^ cells after 4 days of culture (n = 3). ns, not significant * P < .05, ** P < .01, ***P < .001, ****P < .0001.

To further position TRIP13 within the TP53 signaling network, we performed a CRISPR-Cas9 knock-out screen targeting 704 TP53 pathway-associated genes in fetal NUP98::KDM5A^+^ cells, in the presence or absence of *Trip13* deletion (Fig. 4f). This approach enabled systematic identification of genetic modifiers that either mitigated or exacerbated the growth defect induced by *Trip13* loss.

In addition to Trp53 depletion, knock-out of the checkpoint kinase *Chek2* and the pro-apoptotic effector *Bax* were selectively rescued by *Trip13*-deficiency, indicating engagement of a CHEK2–TP53–BAX mitochondrial apoptotic pathway, which was further supported by a rescue of the *Trip13* loss by depletion of the TP53-induced BH3-only protein *Bbc3/Puma* (Fig. 4f), and an upregulation of *Bbc3/Puma* mRNA upon *Trip13* loss (Supplementary Fig. 4d).

Conversely, loss of several negative regulators of TP53 signaling further enhanced the antiproliferative effects of *Trip13* deletion. In particular, knock-out of *Mdm2* and *Bcl-xl/Bcl2l1* synergized with *Trip13* loss, consistent with cooperative derepression of TP53 activity and reinforcement of apoptotic signaling (Fig. 4f). Notably, *Bcl2l1/Bcl-xl* mRNA expression was also increased following *Trip13* deletion, suggesting engagement of compensatory survival feedback mechanisms downstream of TP53 activation (Fig. 4g).

In addition to core TP53 regulators, several genes involved in cell-cycle progression and transcriptional control exhibited enhanced depletion specifically in *Trip13*-deficient cells, including *Cdk1*, *Cdk4*, *Cdk9*, and *Hdac1*, identifying context-specific co-dependencies that emerge under conditions of *Trip13*-loss initiated TP53 signaling (Fig. 4f).

Collectively, these genetic interaction data frames a model in which *TRIP13* restrains TP53 pathway activation through PPM1D/WIP1 and downstream regulators, thereby preventing engagement of a CHEK2–BAX–dependent mitochondrial apoptotic program. Consequently, loss of *TRIP13* unmasks a potent TP53-centered vulnerability in fetal NUP98::KDM5A⁺ leukemia cells that is further modulated by regulators of apoptosis, cell-cycle progression, and transcriptional control.

### Therapeutic Targeting of TRIP13 Reactivates TP53 in NUP98::KDM5A⁺ AML

Our combined proteomic and genomic analyses identified TRIP13 as a critical suppressor of TP53 in fetal NUP98::KDM5A⁺ AML, raising the possibility that pharmacological inhibition of TRIP13 could be therapeutically exploitable [51]. To test this, we treated leukemic cells with DCZ0415, a small-molecule inhibitor of TRIP13. DCZ0415 phenocopied genetic *Trip13* loss and robustly induced apoptosis in NUP98:KDM5A^+^ cells in a dose-dependent manner (mean IC_50_ fetal = 22 µM, adult = 26 µM). NUP98::KDM5A^+^ cells were significantly more sensitive to DCZ0415 than control leukemia models, including FPD-AML and ML-DS, as well as normal adult bone marrow HSPCs (mean IC_50:_ FPD-AML: 50 µM, ML-DS: 35 µM, mBM-HSPCs: 45 µM) (Fig. 5a and Supplementary Fig. 5a-b). Human fetal NUP98::KDM5A⁺ cells exhibited comparable sensitivity (mean IC_50_ = 19 µM), whereas OCI-AML3, Kasumi1 or M07E showed impaired or incomplete responses (IC_50_: OCI-AML3: 32 µM, Kasumi-1/M07e: NM = not measurable; Fig. 5b and Supplementary Fig. 5c).

**Fig. 5.**
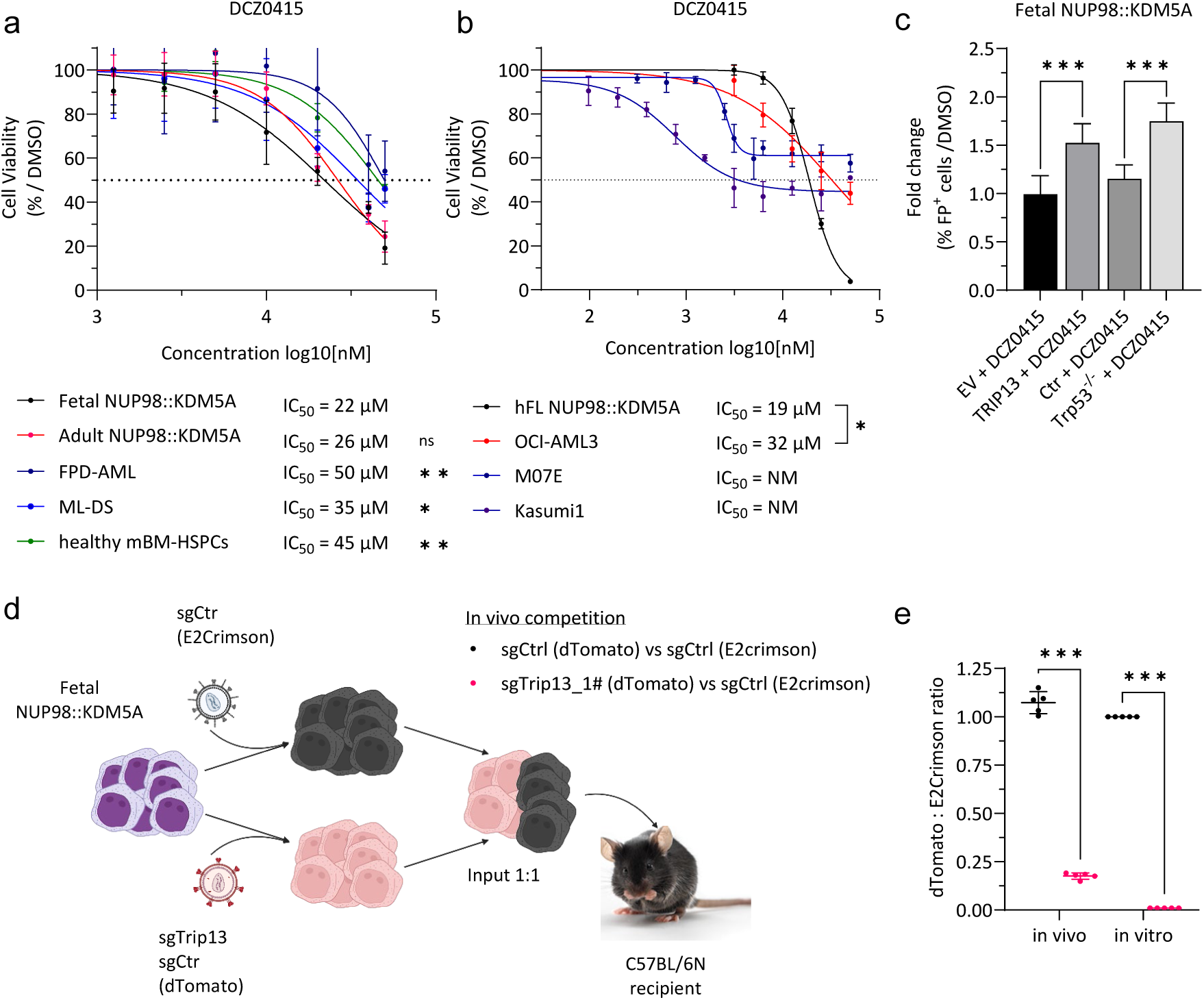
Pharmacological inhibition of the TRIP13-TP53 axis is a therapeutic option for NUP98::KDM5A^+^ pediatric leukemia. a) Dose-response curves for DCZ0415 (TRIP13 inhibitor) in fetal and adult NUP98::KDM5A^+^, ML-DS and FPD-AML models and healthy mBM-HSPCs after 5 days of treatment *in vitro*. The corresponding IC_50_-values are depicted below the graphs. (mean ± s.d., n > 3, 1-way ANOVA related to fetal NUP98::KDM5A^+^ cells) b) Dose-response curves for DCZ0415 (TRIP13 inhibitor) in hFL NUP98::KDM5A^+^ cells and OCI-AML3, M07E and Kasumi1 after 5 days of treatment *in vitro*. The corresponding IC_50_-values are depicted below the graphs. (mean ± s.d., n > 3, 1-way ANOVA related to hFL NUP98::KDM5A^+^ cells) c) Bar graphs showing the percentage of sgRNA-transduced (negative control sgRNA [sgCtr], sgTrp53) and empty vector (EV) or TRIP13-overexpressing fetal NUP98::KDM5A^+^ cells after 7 days of treatment with DCZ0415 (TRIP13 inhibitor) *in vitro*. (mean±s.d., n > 3, 1-way ANOVA) d) Experimental setup for evaluating the genetic ablation of *Trip13* in fetal NUP98::KDM5A^+^ cells *in vivo*. Fetal NUP98::KDM5A^+^ cells (with stable Cas9 expression) were co-transduced with sgTrip13 (dTomato) and sgCtr (dTomato) and mixed 1:1 with sgCtr (E2Crimson), before transplantation into sublethally irradiated recipient mice. e) Dot plot showing the ratio of dTomato^+^ to E2Crimson^+^ cells in the bone marrow of mice euthanized 4 weeks after transplantation (n = 5) and after 4 weeks of culture, normalized to day 0 (n = 3, 2-way ANOVA). ns, not significant * P < .05, ** P < .01, ***P < .001, ****P < .0001.

Importantly, DCZ0415 treatment resulted in activation of the TP53 pathway and cell death, effects that were rescued by enforced TRIP13 expression or by *Trp53* deletion (Fig. 5c, Supplementary Fig. 5d and Supplementary Table 5), confirming on-target activity and TP53-dependent mechanisms of action.

Consistent with the *in vitro* findings, *in vivo* fluorescence-based competitive transplantation assays demonstrated marked depletion of *Trip13*-deficient fetal NUP98::KDM5A⁺ blasts in the bone marrow of recipient mice at the experimental endpoint (Fig. 5d-e).

Together, these data demonstrate that pharmacologic inhibition of TRIP13 reactivates TP53 signaling and selectively compromises the survival of NUP98::KDM5A⁺ leukemic cells, providing functional proof-of-concept that TRIP13 represents a therapeutically targetable vulnerability in this high-risk pediatric AML subtype.

## Discussion

In this study, we adopted a developmental perspective to leukemogenesis to uncover a previously unrecognized vulnerability in pediatric NUP98::KDM5A^+^ AML. Rather than focusing exclusively on oncogenic lesions, we interrogated how fetal versus adult hematopoietic contexts shape leukemic behavior and dependencies. This ontogeny-centered approach revealed a developmentally restricted survival axis centered on TP53 suppression and establishes a conceptual framework that may extend to other malignancies with strong developmental origins.

NUP98::KDM5A-rearranged AML is a rare but highly aggressive leukemia that occurs predominantly in infants and very young children below the age of 2 years [6–8], with clinical and genomic evidence supporting leukemic initiation during embryogenesis [52]. Despite intensive chemotherapy and allogeneic hematopoietic stem cell transplantation (HSCT), outcomes remain poor, relapses are frequent, and treatment-related morbidity is substantial. Achieving deep, measurable residual disease (MRD)–negative remissions before HSCT is crucial for long-term survival, yet current induction regimens are often too toxic or insufficiently effective in this fragile patient population [53]. These challenges underscore the need for biologically informed, mechanism-based therapeutic strategies tailored to the unique features of infant AML.

Ontogeny has emerged as a critical modifier of disease biology in several pediatric leukemias, including CBFA2T3::GLIS2- and KMT2A-rearranged AML [6]. However, its role in NUP98::KDM5A-driven leukemia has not been directly addressed. Using matched fetal and adult HSPC models, we demonstrate that NUP98::KDM5A-driven leukemia arising in a fetal context is more aggressive and retains fetal-specific transcriptional programs [22, 23, 54]. Functional genomic interrogation of fetal-enriched genes identified *TRIP13*, a AAA⁺ ATPase and direct interactor of the TP53 phosphatase PPM1D/WIP1, as a critical dependency in fetal NUP98::KDM5A⁺ cells. Mechanistically, our data show that TRIP13 suppresses TP53 pathway activation and that its genetic or pharmacologic inhibition leads to robust TP53 activation, G₂/M arrest and mitochondrial apoptosis without additional accumulation of DNA damage [45, 46]. These data support a model in which TRIP13 mitigates NUP98::KDM5A-induced DNA stress [55] by modulating TP53 activity, at least in part through its physical interaction with PPM1D, a well-established oncogenic suppressor of TP53 [56]. While the precise molecular mechanism by which TRIP13 regulates PPM1D activity remains to be elucidated, the ATPase-dependent capacity of TRIP13 to remodel protein complexes [57] raises the possibility that it modulates PPM1D conformation, thereby fine-tuning TP53 signaling. Notably, the broader TRIP13 interactome includes multiple TP53 network components, likely accounting for the more pronounced TP53 activation observed upon *TRIP13* loss compared with *PPM1D* deletion alone.

Consistent with this developmental model, our functional screens also identified dependencies on oncofetal regulators previously reported as direct NUP98::KDM5A targets, such as *Igf2bp3* [55, 58], supporting the notion that this fusion sustains leukemic maintenance in part through enforcement of fetal gene expression programs.

Our findings complement prior studies identifying transcriptional and cell-cycle dependencies in NUP98::KDM5A-driven leukemia, including the MEN–KMT2A axis, CDK6, and CDK12 [54, 55, 58]. These dependencies converge on sustained proliferation and transcriptional output, processes inherently associated with replicative stress and DNA damage [59]. In line with these observations, NUP98::KDM5A-expressing cells exhibit a marked baseline DNA damage, reliance on DNA damage response pathways, and heightened sensitivity to CDK12/13 inhibition [55, 60]. Our work adds an essential layer to this model by demonstrating that leukemic survival under such stress conditions requires active control of TP53-mediated apoptosis through a TRIP13-centered mechanism. The overlap between DNA damage-associated genes identified in our screens and those reported to be maintained and upregulated by NUP98::KDM5A during fetal development [55], further supports this integrated model.

Importantly, we provide proof-of-concept that TRIP13 is pharmacologically targetable. The small-molecule inhibitor DCZ0415 faithfully recapitulated the effects of genetic *TRIP13* loss by inducing TP53-dependent apoptosis in NUP98::KDM5A⁺ AML cells, and *Trip13*-depleted cells were selectively impaired *in vivo*. Although DCZ0415 has not yet been evaluated clinically, these data validate TRIP13 as a druggable node and provide a strong rationale for the development of next-generation TRIP13 inhibitors with improved potency and pharmacologic properties suitable for clinical evaluation.

Beyond identifying TRIP13 as a therapeutic target, our TP53-pathway CRISPR screen delineated genetic modifiers that shape the cellular response to TP53 reactivation. Resistance to *TRIP13* loss was conferred by disruption of core apoptotic mediators such as *TP53*, *CHEK2*, *BAX*, and *BBC3/PUMA*, establishing mitochondrial apoptosis as the dominant execution pathway. Conversely, enhanced sensitivity to *TRIP13* loss was observed upon perturbation of cell-cycle and transcriptional regulators, including *CDK1*, *CDK4*, *CDK9*, and *HDAC1*, highlighting conditional co-dependencies emerging under TRIP13-loss induced TP53 signaling. Together, these interactions further underscore the central role of TP53 suppression in sustaining fetal NUP98::KDM5A-driven leukemic survival.

More broadly, our study illustrates how systematic functional interrogation of developmental programs can reveal age- and lineage-specific vulnerabilities that are not apparent from mutation-focused analyses alone. In NUP98::KDM5A⁺ AML, this strategy uncovered a developmentally constrained TRIP13–PPM1D–TP53 survival axis that is essential for leukemic maintenance and therapeutically exploitable. Leveraging developmental biology to expose such context-specific dependencies may represent a powerful paradigm for identifying novel targets in other aggressive pediatric cancers with a clear developmental origin.

## Acknowledgments

The authors thank D. Trono of EPFL (Lausanne, Switzerland), for kindly providing both pMD2.G (plasmid 12259; Addgene) and psPAX2 (plasmid 12260; Addgene); A. Santos (University of Halle-Wittenberg) for assistance with flow cytometry; and K. Huke for assistance with animal experiments. The authors thank C. Hugenberg for assistance and proofreading.

## Funding

L.N.C., M.V., and D.H. were supported by the Frankfurt Foundation for Children with Cancer. This work was supported by the German José Carreras Leukemia Foundation (DJCLS 12 R/2024) and by grants from the parents’ Association “Hilfe für Krebskranke Kinder e.V.” (C^3^OMBAT-AML, L.N.C., J.H.K. and D.H)., the European Research Council Horizon 2020 program (#714226), and the German Research Foundation (DFG; FOR 5433 RNA in Focus; KL-2371/7).

Illustrations (Figures 1a, 2a, 4a and 5d and graphical abstract; https://BioRender.com/3to2ktx, https://BioRender.com/q0hsqlg, https://BioRender.com/qo23x2o, https://BioRender.com/3q2on7c, https://biorender.com/94pt6jg) were created in BioRender.com.

## Contributions

L.N.C. performed experiments, analyzed and interpreted the data, and wrote the manuscript; L.P., H.I., and C.I. performed experiments, analyzed the data, and revised the manuscript; K.S. performed bioinformatics analysis and revised the manuscript; K.M., and L.G. performed experiments; M.V. revised the manuscript; A.S. supervised the analyses and revised the manuscript; D.H. and J.H.K designed the study, analyzed and interpreted the data, wrote the manuscript, and academically drove the project.

## Availability of data and materials

RNA-seq gene expression data have been deposited in the National Center for Biotechnology Information’s Gene Expression Omnibus (accession numbers GSE308953, GSE308954, and GSE308955).

## References

1. Bolouri, H., et al., The molecular landscape of pediatric acute myeloid leukemia reveals recurrent structural alterations and age-specific mutational interactions. Nat Med, 2018. 24(1): p. 103–112.

2. Papaemmanuil, E., et al., Genomic Classification and Prognosis in Acute Myeloid Leukemia. N Engl J Med, 2016. 374(23): p. 2209–2221.

3. Marceau-Renaut, A., et al., Molecular Profiling Defines Distinct Prognostic Subgroups in Childhood AML: A Report From the French ELAM02 Study Group. Hemasphere, 2018. 2(1): p. e31.

4. Reinhardt, D., E. Antoniou, and K. Waack, Pediatric Acute Myeloid Leukemia-Past, Present, and Future. J Clin Med, 2022. 11(3).

5. Elgarten, C.W. and R. Aplenc, Pediatric acute myeloid leukemia: updates on biology, risk stratification, and therapy. Current Opinion in Pediatrics, 2020. 32(1): p. 57–66.

6. Lopez, C.K., et al., Ontogenic Changes in Hematopoietic Hierarchy Determine Pediatric Specificity and Disease Phenotype in Fusion Oncogene-Driven Myeloid Leukemia. Cancer Discov, 2019. 9(12): p. 1736–1753.

7. Okeyo-Owuor, T., et al., The efficiency of murine MLL-ENL-driven leukemia initiation changes with age and peaks during neonatal development. Blood Adv, 2019. 3(15): p. 2388–2399.

8. Calvi, L.M. and D.C. Link, The hematopoietic stem cell niche in homeostasis and disease. Blood, 2015. 126(22): p. 2443–51.

9. Mack, R., et al., The Fetal-to-Adult Hematopoietic Stem Cell Transition and its Role in Childhood Hematopoietic Malignancies. Stem Cell Rev Rep, 2021. 17(6): p. 2059–2080.

10. de Smith, A.J. and L.G. Spector, In Utero Origins of Acute Leukemia in Children. Biomedicines, 2024. 12(1).

11. Gruber, T.A. and J.R. Downing, The biology of pediatric acute megakaryoblastic leukemia. Blood, 2015. 126(8): p. 943–9.

12. de Rooij, J.D., et al., Pediatric non-Down syndrome acute megakaryoblastic leukemia is characterized by distinct genomic subsets with varying outcomes. Nat Genet, 2017. 49(3): p. 451–456.

13. de Rooij, J.D., et al., Recurrent abnormalities can be used for risk group stratification in pediatric AMKL: a retrospective intergroup study. Blood, 2016. 127(26): p. 3424–30.

14. Gough, S.M., C.I. Slape, and P.D. Aplan, NUP98 gene fusions and hematopoietic malignancies: common themes and new biologic insights. Blood, 2011. 118(24): p. 6247–57.

15. Bertrums, E.J.M., et al., Comprehensive molecular and clinical characterization of NUP98 fusions in pediatric acute myeloid leukemia. Haematologica, 2023.

16. Gough, S.M., et al., NUP98-PHF23 is a chromatin-modifying oncoprotein that causes a wide array of leukemias sensitive to inhibition of PHD histone reader function. Cancer Discov, 2014. 4(5): p. 564–77.

17. Wang, G.G., et al., Haematopoietic malignancies caused by dysregulation of a chromatin-binding PHD finger. Nature, 2009. 459(7248): p. 847–51.

18. Wang, G.G., et al., NUP98-NSD1 links H3K36 methylation to Hox-A gene activation and leukaemogenesis. Nat Cell Biol, 2007. 9(7): p. 804–12.

19. Bowie, M.B., et al., Identification of a new intrinsically timed developmental checkpoint that reprograms key hematopoietic stem cell properties. Proc Natl Acad Sci U S A, 2007. 104(14): p. 5878–82.

20. Popescu, D.M., et al., Decoding human fetal liver haematopoiesis. Nature, 2019. 574(7778): p. 365–371.

21. Ranzoni, A.M., et al., Integrative Single-Cell RNA-Seq and ATAC-Seq Analysis of Human Developmental Hematopoiesis. Cell Stem Cell, 2021. 28(3): p. 472–487.e7.

22. Roy, A., et al., Transitions in lineage specification and gene regulatory networks in hematopoietic stem/progenitor cells over human development. Cell Rep, 2021. 36(11): p. 109698.

23. Ivanovs, A., et al., Human haematopoietic stem cell development: from the embryo to the dish. Development, 2017. 144(13): p. 2323–2337.

24. Gialesaki, S., et al., RUNX1 isoform disequilibrium promotes the development of trisomy 21–associated myeloid leukemia. Blood, 2023. 141(10): p. 1105–1118.

25. Alejo-Valle, O., et al., The megakaryocytic transcription factor ARID3A suppresses leukemia pathogenesis. Blood, 2022. 139(5): p. 651–665.

26. Emmrich, S., et al., miR-99a/100∼125b tricistrons regulate hematopoietic stem and progenitor cell homeostasis by shifting the balance between TGFβ and Wnt signaling. Genes Dev, 2014. 28(8): p. 858–74.

27. Bhayadia, R., et al., Endogenous Tumor Suppressor microRNA-193b: Therapeutic and Prognostic Value in Acute Myeloid Leukemia. J Clin Oncol, 2018. 36(10): p. 1007–1016.

28. Labuhn, M., et al., Refined sgRNA efficacy prediction improves large- and small-scale CRISPR-Cas9 applications. Nucleic Acids Res, 2018. 46(3): p. 1375–1385.

29. Reimer, J., et al., CRISPR-Cas9-induced t(11;19)/MLL-ENL translocations initiate leukemia in human hematopoietic progenitor cells in vivo. Haematologica, 2017. 102(9): p. 1558–1566.

30. de Rooij, J.D., et al., NUP98/JARID1A is a novel recurrent abnormality in pediatric acute megakaryoblastic leukemia with a distinct HOX gene expression pattern. Leukemia, 2013. 27(12): p. 2280–8.

31. Labuhn, M., et al., Mechanisms of Progression of Myeloid Preleukemia to Transformed Myeloid Leukemia in Children with Down Syndrome. Cancer Cell, 2019. 36(2): p. 123–138.e10.

32. Li, W., et al., MAGeCK enables robust identification of essential genes from genome-scale CRISPR/Cas9 knockout screens. Genome Biol, 2014. 15(12): p. 554.

33. Ianevski, A., A.K. Giri, and T. Aittokallio, SynergyFinder 3.0: an interactive analysis and consensus interpretation of multi-drug synergies across multiple samples. Nucleic Acids Research, 2022. 50(W1): p. W739–W743.

34. Chen, S., et al., fastp: an ultra-fast all-in-one FASTQ preprocessor. Bioinformatics, 2018. 34(17): p. i884–i890.

35. Spinozzi, G., et al., ARPIR: automatic RNA-Seq pipelines with interactive report. BMC Bioinformatics, 2020. 21(Suppl 19): p. 574.

36. Love, M.I., W. Huber, and S. Anders, Moderated estimation of fold change and dispersion for RNA-seq data with DESeq2. Genome Biol, 2014. 15(12): p. 550.

37. Benjamini, Y. and Y. Hochberg, Controlling the False Discovery Rate: A Practical and Powerful Approach to Multiple Testing. Journal of the Royal Statistical Society: Series B (Methodological), 1995. 57(1): p. 289–300.

38. Subramanian, A., et al., Gene set enrichment analysis: A knowledge-based approach for interpreting genome-wide expression profiles. Proceedings of the National Academy of Sciences, 2005. 102(43): p. 15545–15550.

39. Mootha, V.K., et al., PGC-1α-responsive genes involved in oxidative phosphorylation are coordinately downregulated in human diabetes. Nature Genetics, 2003. 34(3): p. 267–273.

40. Harrison, P.W., et al., Ensembl 2024. Nucleic Acids Research, 2023. 52(D1): p. D891–D899.

41. Noort, S., et al., The clinical and biological characteristics of NUP98-KDM5A in pediatric acute myeloid leukemia. Haematologica, 2021. 106(2): p. 630–634.

42. Schwarzer, A., et al., The non-coding RNA landscape of human hematopoiesis and leukemia. Nat Commun, 2017. 8(1): p. 218.

43. Förster, A., et al., Beyond Pathogenic RUNX1 Germline Variants: The Spectrum of Somatic Alterations in RUNX1-Familial Platelet Disorder with Predisposition to Hematologic Malignancies. Cancers (Basel), 2022. 14(14).

44. Grimm, J., D. Heckl, and J.H. Klusmann, Molecular Mechanisms of the Genetic Predisposition to Acute Megakaryoblastic Leukemia in Infants With Down Syndrome. Front Oncol, 2021. 11: p. 636633.

45. Vader, G., Pch2TRIP13: controlling cell division through regulation of HORMA domains. Chromosoma, 2015. 124(3): p. 333–339.

46. Banerjee, R., et al., TRIP13 promotes error-prone nonhomologous end joining and induces chemoresistance in head and neck cancer. Nat Commun, 2014. 5: p. 4527.

47. Jeong, H., et al., TRIP13 Participates in Immediate-Early Sensing of DNA Strand Breaks and ATM Signaling Amplification through MRE11. Cells, 2022. 11(24).

48. Sullivan, K.D., et al., Mechanisms of transcriptional regulation by p53. Cell Death & Differentiation, 2018. 25(1): p. 133–143.

49. Kruse, J.-P. and W. Gu, Modes of p53 Regulation. Cell, 2009. 137(4): p. 609–622.

50. Andrysik, Z., et al., PPM1D suppresses p53-dependent transactivation and cell death by inhibiting the Integrated Stress Response. Nat Commun, 2022. 13(1): p. 7400.

51. Wang, Y., et al., A Small-Molecule Inhibitor Targeting TRIP13 Suppresses Multiple Myeloma Progression. Cancer Res, 2020. 80(3): p. 536–548.

52. Cazzola, A., et al., Prenatal Origin of Pediatric Leukemia: Lessons From Hematopoietic Development. Front Cell Dev Biol, 2020. 8: p. 618164.

53. Rettinger, E., et al., The hallmarks of hematopoietic stem cell transplantation for pediatric acute myeloid leukemia. Leukemia, 2025.

54. Heikamp, E.B., et al., The menin-MLL1 interaction is a molecular dependency in NUP98-rearranged AML. Blood, 2022. 139(6): p. 894–906.

55. Troester, S., et al., Transcriptional and epigenetic rewiring by the NUP98::KDM5A fusion oncoprotein directly activates CDK12. Nat Commun, 2025. 16(1): p. 4656.

56. Ogasawara, S., et al., Novel inhibitors targeting PPM1D phosphatase potently suppress cancer cell proliferation. Bioorg Med Chem, 2015. 23(19): p. 6246–9.

57. Mapelli, M. and A. Musacchio, MAD contortions: conformational dimerization boosts spindle checkpoint signaling. Curr Opin Struct Biol, 2007. 17(6): p. 716–25.

58. Schmoellerl, J., et al., CDK6 is an essential direct target of NUP98 fusion proteins in acute myeloid leukemia. Blood, 2020. 136(4): p. 387–400.

59. Shimura, T., et al., Cyclin D1 overexpression perturbs DNA replication and induces replication-associated DNA double-strand breaks in acquired radioresistant cells. Cell Cycle, 2013. 12(5): p. 773–82.

60. Domingo-Reinés, J., et al., The pediatric leukemia oncoprotein NUP98-KDM5A induces genomic instability that may facilitate malignant transformation. Cell Death Dis, 2023. 14(6): p. 357.

