## Supplementary data for "Activation of the TP53 pathway is a therapeutic vulnerability in NUP98::KDM5A^+^ pediatric AML"

#### Supplementary Figures

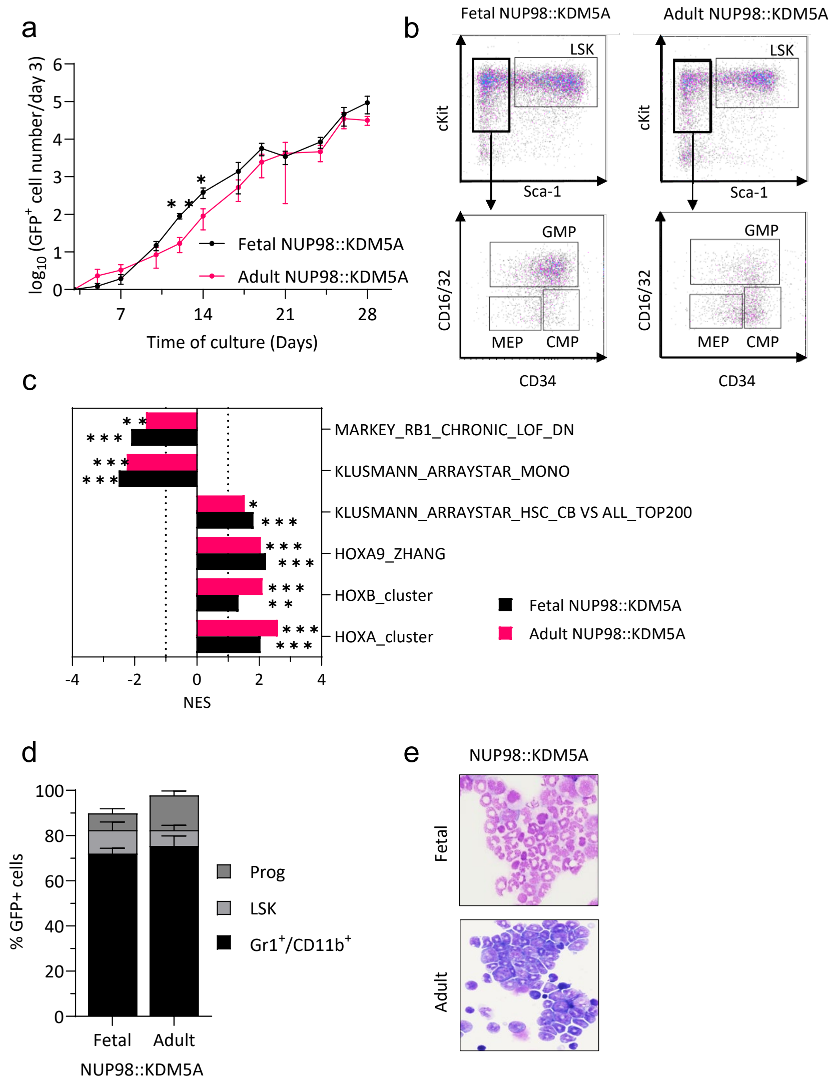


**Supplementary Fig. 1. NUP98::KDM5A fusion oncoprotein induces a more aggressive disease in fetal cells.**

a) $Log_{10}$ $GFP^+$ cell number of fetal and adult NUP98::KDM5A$^+$ maintained in liquid culture. Data from 1 representative experiment performed in replicates (n = 3) are shown as mean s.d. (t-test).
b) Representative flow cytometry analysis of fetal and adult NUP98::KDM5A$^+$ cells after 4 weeks of culture.
c) Bar graphs showing the normalized enrichment scores (NES) of significantly upregulated or downregulated gene sets in fetal and adult NUP98::KDM5A$^+$ cells compared to the respective control (EV).
d) Bar graph showing the percentage of LSK (Lin$^-$. c-kit$^+$ and Sca1$^+$). Progenitor (Lin$^-$. c-kit$^+$ and Sca1$^-$) and myeloid (Gr1$^+$ and CD11b$^+$) population in leukemic blasts isolated from bone marrow of mice transplanted with fetal and adult NUP98-KDM5A$^+$ cells.
e) Representative bone marrow cytospin images (x40 original magnification) of C57BL/6N mice transplanted with fetal and adult NUP98-KDM5A$^+$ cells.

ns. not significant * P < .05. ** P < .01. ***P < .001. ****P < .0001.

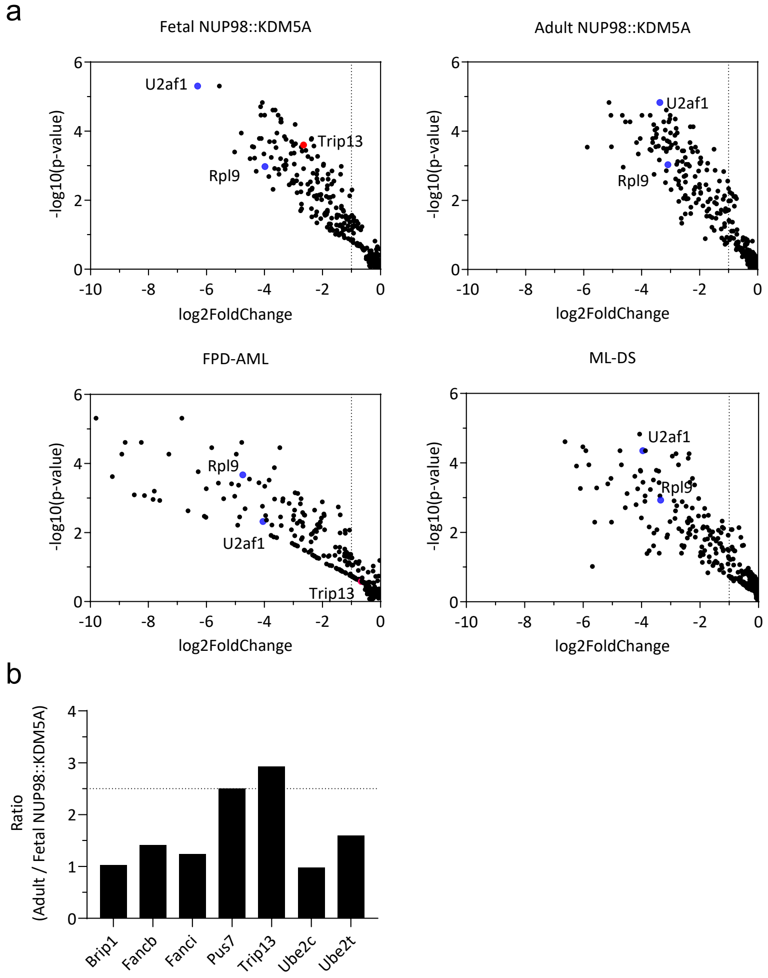


**Supplementary Fig. 2. CRISPR-Cas9 knock-out screening reveals *TRIP13* dependency in fetal NUP98::KDM5A$^{+}$ AML.**

a) Dot plots showing the log2 fold change and –log10(p-value) of the fetal enriched gene targeted by sgRNAs in fetal (top-left), adult (top-right) NUP98::KDM5A, FPD-AML (bottom-left) and ML-DS (bottom-right) models. Blue dots represent positive controls (*Rpl9* and *U2af1*). Red dots represent selected candidate gene *Trip13*. (n =2)
b) Bar plot showing the ratio between adult and fetal NUP98::KDM5A models of the seven top candidates of fetal NUP98::KDM5A model.

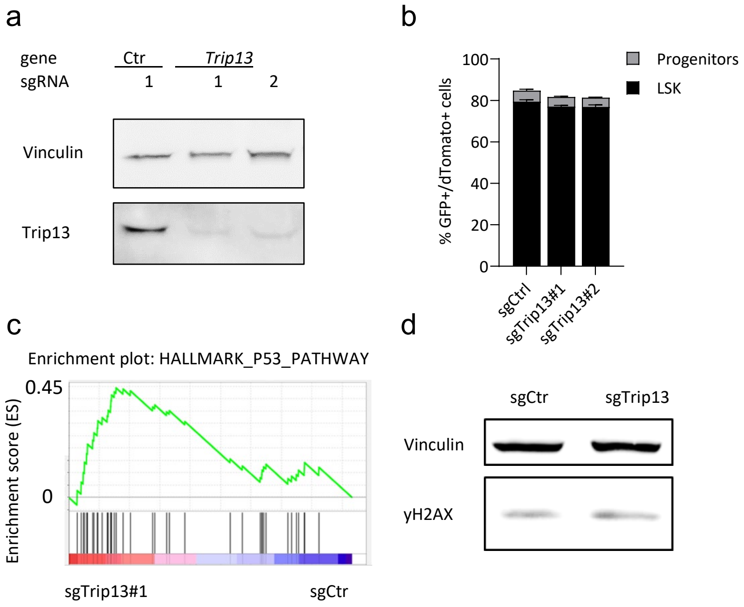


**Supplementary Fig. 3. *Trip13* loss promotes activation of the TP53 pathway and arrests fetal NUP98::KDM5A$^+$ cell proliferation *in vitro*.**

a) Western blot showing Trip13 protein levels in sorted sgRNA-transduced (negative control sgRNA [Luc]. sgTrip13#1-2) fetal NUP98::KDM5A$^+$ cells 4 days after transduction. Endogenous Vinculin was used as loading control.
b) Bar graph showing the percentage of LSK (Lineage negative. c-kit$^+$ and Sca1$^+$) and progenitor (Lineage negative. c-kit$^+$. Sca1$^-$) subsets in sgRNA-transduced (negative control sgRNA [sgCtr]. sgTrip13#1-2) fetal NUP98::KDM5A$^+$ cells after 5 days of culture (mean ± s.d.. n > 3 per sgRNA. 2-way ANOVA).
c) Gene Set Enrichment Analysis (GSEA) plot showing deregulated genes involved in TP53 pathway in *Trip13*-depleted fetal NUP98::KDM5A$^+$ cells compared to control.
d) Western blot showing yH2AX protein level in sorted sgRNA-transduced (negative control sgRNA [Luc]. sgTrip13) fetal NUP98::KDM5A$^+$ cells 4 days after transduction. Endogenous Vinculin was used as loading control.

ns. not significant * P < .05. ** P < .01. ***P < .001. ****P < .0001.

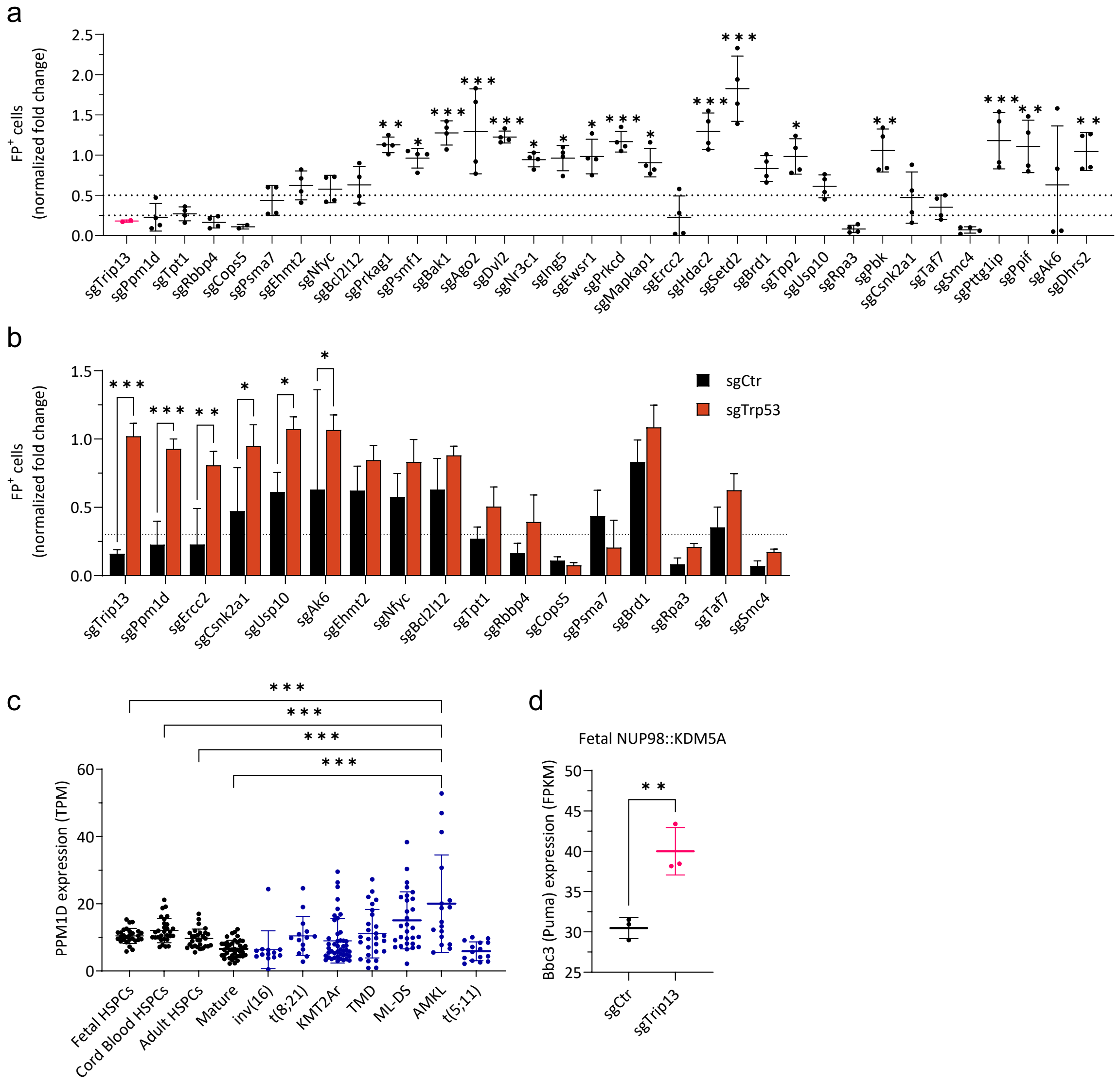


**Supplementary Fig. 4. TRIP13 interacts with PPM1D/WIP1 to repress the TP53 pathway.**

a) Dot plot showing the percentage of sgRNA-transduced fetal NUP98::KDM5A$^{+}$ cells after 14 days of culture, normalized to day 2 ( mean±s.d., n > 3 per sgRNA, 1-way ANOVA).
b) Bar plot showing the percentage of sgRNA-transduced fetal NUP98::KDM5A$^{+}$ cells in combination with negative control sgRNA [sgCtr] or sgTrp53, after 14 days of culture, normalized to day 2 ( mean±s.d., n > 3 per sgRNA. 1-way ANOVA).
c) Dot plot showing the expression of *PPM1D* in subgroups of HSPCs and pediatric AML patients from the HemAtlas dataset.
d) Dot plot showing the expression of *Bbc3/Puma* in control or *Trip13*-depleted fetal NUP98::KDM5A$^{+}$ cells after 4 days of culture (n = 3)

ns. not significant * P < .05. ** P < .01. ***P < .001. ****P < .0001.

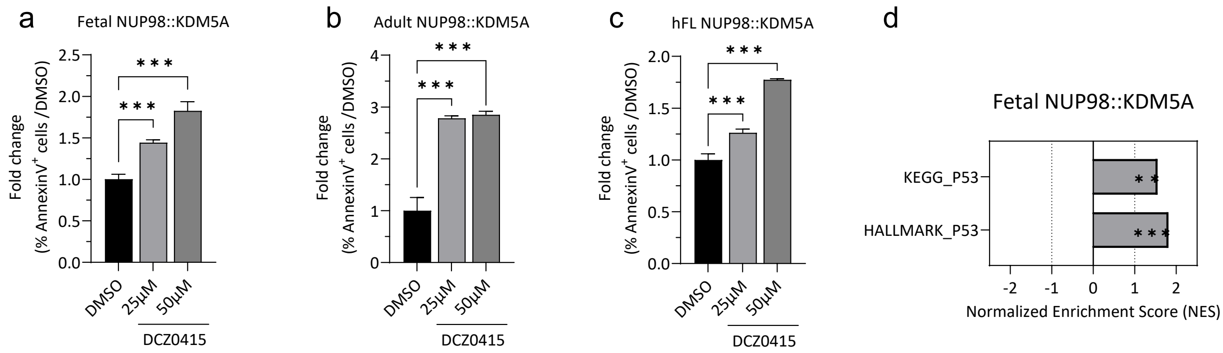


**Supplementary Fig. 5. Pharmacological inhibition of TRIP13-TP53 axis is a therapeutic option for NUP98::KDM5A$^+$ pediatric AML.**

a) Bar graphs showing the percentage of Annexin-V$^+$ cells after treatment with the indicated doses of DCZ0415 compared to dimethyl sulfoxide control (DMSO) in fetal NUP98::KDM5A$^+$ cells after 5 days of treatment *in vitro*. (mean±s.d., n > 3. 1-way ANOVA)
b) Bar graphs showing the percentage of Annexin-V$^+$ cells after treatment with the indicated doses of DCZ0415 compared to dimethyl sulfoxide control (DMSO) in adult NUP98::KDM5A$^+$ cells after 5 days of treatment *in vitro*. (mean±s.d., n > 3. 1-way ANOVA
c) Bar graphs showing the percentage of Annexin-V$^+$ cells after treatment with the indicated doses of DCZ0415 compared to dimethyl sulfoxide (DMSO) control in hFL NUP98::KDM5A$^+$ cells after 5 days of treatment *in vitro*. (mean±s.d., n > 3. 1-way ANOVA).
d) Bar graphs showing the normalized enrichment scores (NES) of significantly upregulated or downregulated gene sets in fetal NUP98::KDM5A$^+$ cells after 24 hours DCZ0415 treatment compared dimethyl sulfoxide (DMSO) control.

ns. not significant * P < .05. ** P < .01. ***P < .001.

### Supplementary Tables

**Supplementary Table 1.** List of reagents and resources

| Reagents | Company or source | Identifier |
|---|---|---|
| **Antibodies** | | |
| Anti-human CD41-PeCy7; clone P2 | Beckman Coulter | Cat# 6607115. RRID: AB_2800448 |
| Anti-human CD42b-APC ; clone HIP1 | BD Biosciences | Cat# 551061. RRID:AB_398486 |
| Anti-mouse Ly-6G/Ly-6C-APC ; clone RB6-8C5 | Biolegend | Cat# 108412. RRID:AB_313377 |
| Anti-mouse CD41-APC-Cy7; clone MWReg30 | Biolegend | Cat# 133928. RRID:AB_2572132 |
| Anti-mouse CD117 (c-kit) APC/Cyanine7; clone 2B8 | BD biosciences | Cat# 105826. RRID:AB_1626278 |
| Anti-mouse/rat CD42d APC; clone 1C2 | BD biosciences | Cat# 148505. RRID:AB_2564601 |
| Anti-mouse/rat CD42d PerCP/Cyanine5.5; clone 1C2 | BD biosciences | Cat# 148507. RRID:AB_2564603 |
| Anti-mouse CD117 (c-kit) APC/Cyanine7; clone 2B8 | BD biosciences | Cat# 105826. RRID:AB_1626278 |

| | | |
|---|---|---|
| Anti-mouse CD34 PerCP/Cyanine5.5; clone HM34 | BD biosciences | Cat# 128607. RRID:AB_1279222 |
| Anti-mouse CD71 PerCP/Cyanine5.5; clone CY1G4 | BD biosciences | Cat# 567256. RRID:AB_2916518 |
| Anti-mouse Ly-6A/E PE/Cy7; clone D7 | BD biosciences | Cat# 561021. RRID:AB_2034021 |
| Anti-CD11b PE/Cy7; clone M1/70 | BD biosciences | Cat# 552850. RRID:AB_394491 |
| Anti-Human CD38 PE/Cy7; clone HB7 | BD biosciences | Cat# 335825. RRID:AB_2868688 |
| Anti-Human CD90 APC; clone 5e10 | BD biosciences | Cat# 559869. RRID:AB_398677 |
| Anti-Human CD34 PE/Cy7; clone 581 | Beckman Coulter | Cat# A21691 |
| Anti-Human CD235a APC-A750; clone 11E4B-7-6 (KC16) | Beckman Coulter | Cat# A89314 |
| Anti-Human CD33 PerCP/Cyanine5.5; clone D3HL60.251 | Beckman Coulter | Cat# A70198 |
| Anti-Human CD117 APC-A750; clone 104D2D1 | Beckman Coulter | Cat# B92450 |
| Anti-mouse TER-119 APC/Cyanine7; clone TER-119 | BD biosciences | Cat# 560509. RRID:AB_1645230 |
| Anti-mouse CD16/32 APC/Cyanine7; clone S17011E | BD biosciences | Cat# 156612. RRID:AB_2800710 |
| Anti-mouse Ly-6G/Ly-6C PerCP/Cyanine5.5; clone RB6-8C5 | BD Biosciences | Cat# 552093. RRID:AB_394334 |
| Anti-mouse CD117-APC | BD Biosciences | Cat# 553356. RRID:AB_398536 |
| Anti-mouse TER-119-Biotin. clone TER-119 | BD Biosciences | Cat#553672. RRID:AB_394985 |
| Anti-HA-Tag (C29F4) Rabbit mAb | Cell signaling | Cat# 3724. RRID:AB_1549585 |
| Anti-Vinculin (E1E9V) XP® Rabbit | abcam | Cat# 13901. RRID:AB_2728768 |
| TRIP13 Antibody (C-4) | Santa Cruz | Cat# sc-514314 |
| Phospho-Histone H2A.X (Ser139) (20E3) Rabbit mAb | Cell Signaling | Cat# 9718T |
| p53 (D2H9O) Rabbit mAb | Cell Signaling | Cat# 32532S. RRID:AB_2757821 |
| Recombinant Anti-PPM1D/WIP1 antibody [EPR22960-39] (ab234439) | abcam | Cat# ab234439 |
| Vinculin (E1E9V) XP Rabbit mAb | Cell Signaling | Cat# 13901S. RRID:AB_2728768 |
| Anti-HA tag antibody [HA.C5] | Cell Signaling | Cat# ab18181. RRID:AB_444303 |
| Rabbit Anti-mouse IgG H&L | abcam | Cat# ab46540. RRID:AB_2614925 |

| | | |
|---|---|---|
| Lineage cocktail-Pacific Blue; clone 17A2; RB6-8C5; RA3-6B2; Ter-119; M1/70 | BioLegend | Cat# 133310; RRID:AB_11150779 |
| Pierce™ anti-HA magnetic beads | ThermoFisher Scientific | Cat# 88837. RRID:AB_2861399 |
| APC-Annexin V | BD Biosciences | Cat# 550475. RRID:AB_2868885 |
| PE/Cyanine7 anti-BrdU Antibody Clone 3D4 | Biolegend | Cat# 364118. RRID:AB_2814319 |
| **Cytokines. peptides and chemicals** | | |
| Lipofectamine 2000 Transfection Reagent | Thermo Fisher Scientific | Cat# 11668027 |
| Recombinant Human TPO | Peprotech | Cat# 300-18 |
| Recombinant Human SCF | Peprotech | Cat# 300-07 |
| Recombinant Human IL-3 | Peprotech | Cat# 200-03 |
| Recombinant Human Flt3-Ligand | Peprotech | Cat# 300-19 |
| Recombinant Human IL-6 | Peprotech | Cat# 200-06 |
| Recombinant Murine TPO | Peprotech | Cat# 315-14 |
| Recombinant Murine SCF | Peprotech | Cat# 250-03 |
| cOmplete™. Mini Protease Inhibitor Cocktail | Sigma Aldrich | Cat# 04693124001 |
| StemRegenin 1 | STEMCELL Technologies | Cat# 72344 |
| UM171 | STEMCELL Technologies | Cat# 72914 |
| RetroNectin® GMP grade | Takara Bio | Cat# T202 |
| Polybrene | Sigma Aldrich | Cat# TR-1003-G |
| Erythrocyte Lysis buffer | c-c-Pro | Cat# PL-29-L |
| Doxycycline | Sigma Aldrich | Cat# D9891-1G |
| RIPA Lysis and Extraction Buffer | ThermoFisher Scientific | Cat# 89900 |
| 4x Laemmli Sample Buffer | Biorad | Cat# 1610747 |
| 2-mercaptoethanol | Sigma-Aldrich | Cat# M7154 |
| Protease Inhibitor Cocktail (PIC) | Sigma-Aldrich | Cat# P8340 |
| PMSF Protease Inhibitor | ThermoFisher Scientific | Cat# 36978 |
| Sodium-orthovanadate | Merckmillipore | Cat# 567540 |
| CellTiter-Glo Luminescent Cell viability Assay | Promega | Cat# G7571 |
| DTT | CarlRoth | Cat# 6908.1 |
| UREA | Merckmillipore | Cat# 108487 |
| DCZ0415 | MedChemExpress | Cat# HY-15676 |
| **Comercially available kits** | | |

| | | |
|---|---|---|
| GeneJET Gel Extraction Kit | Thermo Fisher Scientific | Cat# K0691 |
| QIAGEN Plasmid Maxi Kit | Qiagen | Cat# 12163 |
| EasySep Mouse Streptavidin RapidSpheres Isolation Kit | Stemcell Technologies | Cat# 19860 |
| QIAamp DNA Blood Mini Kit | Qiagen | Cat# 51104 |
| Quick RNA Microprep Kit | Zymo Research | Cat# R1050 |
| EasySep Human CD34 Positive Selection Kit | Stemcell Technologies | Cat# 17856 |
| NEBNext® High-Fidelity 2X PCR Master Mix | New England Biolabs | Cat# M0541L |
| Amersham™ ECL Prime Western Blotting Detection Reagent | Thermo Fisher Scientific | Cat# 12316992 |
| APC Annexin V Apoptosis Detection Kit II | BD Biosciences | Cat# 560209 |
| GeneJET Plasmid Miniprep Kit | ThermoFisher Scientific | Cat# K0502 |
| Polyethylenimine. Linear. MW 25000. Transfection Grade (PEI 25K™) | Polysciences | Cat# 23966 |
| Biotin Rat Anti-Mouse TER-119/Erythroid Cells | BD | Cat# 553672 |
| CD34 MicroBead Kit UltraPure. human | Miltenyi Biotech | Cat# 130-100-453 |
| APC BrdU Flow Kit | BD Biosciences | Cat# 552598 |
| **Deposited data** | | |
| Raw sequencing data (RNA-seq profiling of NUP98::KDM5A and control (EV) transduced mFL-HSPCs and mBM-HSPCs at day 3 and 21 of culture) | Gene Expression Omnibus: https://www.ncbi.nlm.nih.gov/geo/ | study accession number: GSE308953 |
| Raw sequencing data (RNA-seq-based gene expression analysis of sgRNA-transduced (negative control sgRNA [sgCtr], sgTrip13) fetal NUP98::KDM5A+ cells was performed at day 4 post-transduction) | Gene Expression Omnibus: https://www.ncbi.nlm.nih.gov/geo/ | study accession number: GSE308954 |
| Raw sequencing data (RNA-seq-based gene expression analysis of fetal NUP98::KDM5A+ cells treated for 24 hours with DCZ0415 or DMSO control) | Gene Expression Omnibus: https://www.ncbi.nlm.nih.gov/geo/ | study accession number: GSE308955 |
| **Recombinant DNA** | | |

| | | |
|---|---|---|
| psPAX2 | N/A | Addgene Plasmid #12260; RRID:Addgene_12260 |
| pMD2.G | N/A | Addgene Plasmid #12259; RRID:Addgene_12259 |
| SGL40C.EFS.dTomato | Labuhn et al., 2018 | Addgene Plasmid #89395; RRID:Addgene_89395 |
| SGL40C.EFS.E2Crimson | Labuhn et al., 2018 | Addgene Plasmid #100894; RRID:Addgene_100894 |
| SIN40C.SFFV.GFP.IRES.MCS | This paper | |
| SIN40C.SFFV.MCS.IRES.dTomato | Alejo-Valle et al., 2021 | Addgene Plasmid #169279 |
| SIN40C.TRE.MCS.IRES.dTomato.PGK.sfGFP.P2A.Tet3G | Alejo-Valle et al., 2021 | Addgene Plasmid #169283 |
| SIN40C.SFFV.GFP.IRES.NUP98::KDM5A | This paper | |
| SIN40C.SFFV.TRIP13.IRES.dTomato | This paper | |
| **Software** | | |
| Graphpad Prism version 8 | Graphpad Software | https://www.graphpad.com/scientific-software/prism/ RRID:SCR_002798 |
| R software 3.6.1 | R software | https://www.r-project.org/about.html; RRID:SCR_001905 |
| Kaluza 1.5 | Beckman Coulter | https://www.beckman.de/flow-cytometry/software/kaluza<br>RRID:SCR_016182 |
| Snapgene 5.2 | Insightful Science | https://www.snapgene.com/<br>RRID:SCR_015052 |
| SequestHT | Eng et al., 1994 | http://fields.scripps.edu/yates/wp/?page_id=17 |
| GSEA version 3.0 | Subramanian et al., 2005 | https://software.broadinstitute.org/gsea/index.jsp; RRID:SCR_003199 |
| Proteome Discoverer 2.4 | Thermo Fisher Scientific | Cat#OPTON-30945; RRID:SCR_014477 |
| CytExpert Software | Beckman Coulter | https://www.beckman.com/flow-cytometry/instruments/cytoflex/software; RRID:SCR_017217 |
| MAGeCK | Li et al., 2014 | https://sourceforge.net/p/mageck/wiki/Home/ |
| BZ-II Viewer | Keyence | https://www.keyence.com/ |
| BZ-II Analyzer | Keyence | https://www.keyence.com/ |

| | | |
|---|---|---|
| Cytoscape | Shannon et al., 2003 | http://cytoscape.org; RRID:SCR_003032 |
| STRING | Doncheva et al., 2019 | http://string.embl.de/RRID:SCR_005223 |
| Ensembl | Aken et al., 2017 | http://www.ensembl.org/index.html; RRID:SCR_2344 |

**Supplementary Table 2.** List of oligonucleotides

| sgRNAs Name | FW_Sequence | REV_Sequence |
|---|---|---|
| sgTrp53.4 | CACCGAGGAGCTCCTGACACTCGGA | AAACTCCGAGTGTCAGGAGCTCCTC |
| sgTrip13#1 | CACCGGATAGCCTCGTGTATGATG | AAACCATCATACACGAGGCTATCC |
| sgTrip13#2 | CACCGCTACTCGGATAGCATCTGA | AAACTCAGATGCTATCCGAGTAGC |
| sgTrip13#3 | AAACTCAGATGCTATCCGAGTAGC | AAACTCAGGTAAATTCCTAGTTC |
| sgIng5#1 | CACCgCCTGCACTCACCATCTCGT | AAACACGAGATGGTGAGTGCAGGc |
| sgIng5#2 | CACCGAAAGCAGAGATCGACATCC | AAACGGATGTCGATCTCTGCTTTC |
| sgEhmt2#1 | CACCGTGAGCTACACGAAAGTCG | AAACCGACTTTCGTGTAGCTCAC |
| sgEhmt2#2 | CACCGAGTGATGCGGCCCCGACA | AAACTGTCGGGGCCGCATCACTC |
| sgNr3c1#1 | CACCgATTATGGGGTGCTGACGTG | AAACCACGTCAGCACCCCATAATc |
| sgNr3c1#2 | CACCgTGTCCATGGGACTGTATAT | AAACATATACAGTCCCATGGACAc |
| sgNfyc#1 | CACCGAAATCCGAAACTTAACAG | AAACCTGTTAAGTTTCGGATTTC |
| sgNfyc#2 | CACCgAGTACTGGACAGGCTCCGC | AAACGCGGAGCCTGTCCAGTACTc |
| sgBcl2l12#1 | CACCgAGGCTCGGAACCATAGCAG | AAACCTGCTATGGTTCCGAGCCTc |
| sgBcl2l12#2 | CACCgCCCTGTCCCAACTCCACCC | AAACGGGTGGAGTTGGGACAGGGc |
| sgPpm1d#1 | CACCgTAGCTCCACAAGTCACCTA | AAACTAGGTGACTTGTGGAGCTAc |
| sgPpm1d#2 | CACCGAATGGCCAAAGACTATGAC | AAACGTCATAGTCTTTGGCCATTC |
| sgPpif#1 | CACCGATGTCGTGCCAAAGACTGC | AAACGCAGTCTTTGGCACGACATC |
| sgPpif#2 | CACCgCGCTCGTGTACTTGGACGT | AAACACGTCCAAGTACACGAGCGc |
| sgBak1#1 | CACCGAACTCTGTGTCGTAGCGC | AAACGCGCTACGACACAGAGTTC |
| sgBak1#2 | CACCgTGGTACCTGGAGGCGATCT | AAACAGATCGCCTCCAGGTACCAc |
| sgAgo2#1 | CACCgCTGTCCGACTTGTAAACAC | AAACGTGTTTACAAGTCGGACAGc |
| sgAgo2#2 | CACCGATACCTGTTCACTCTCCGA | AAACTCGGAGAGTGAACAGGTATC |
| sgAk6#1 | CACCGTTATACGACGGCTACGATG | AAACCATCGTAGCCGTCGTATAAC |
| sgAk6#2 | CACCgCTATATGAAACCAGCGTTC | AAACGAACGCTGGTTTCATATAGc |
| sgDhrs2#1 | CACCgACTGGTACATAAGCCACTC | AAACGAGTGGCTTATGTACCAGTc |
| sgDhrs2#2 | CACCGGGAGCCAGTGAACAGATC | AAACGATCTGTTCACTGGCTCCC |
| sgDvl2#1 | CACCgCGAATCTGTCGTATCACTG | AAACCAGTGATACGACAGATTCGC |

| | | |
|---|---|---|
| sgDvl2#2 | CACCGAATACCTAGAAAGGCGTTT | AAACAAACGCCTTTCTAGGTATTC |
| sgTpt1#1 | CACCgCAGCCCGTCCGCGATCTCC | AAACGGAGATCGCGGACGGGCTGc |
| sgTpt1#2 | CACCgCCGATGAGCGAGTCATCGA | AAACTCGATGACTCGCTCATCGGc |
| sgRbbp4#1 | CACCGTAAGTGCCCACTGAGATT | AAACAATCTCAGTGGGCACTTAC |
| sgRbbp4#2 | CACCGCCACTGGGCAGTTAAGCT | AAACAGCTTAACTGCCCAGTGGC |
| sgPrkag1#1 | CACCGTGTCAAATACCACCAACT | AAACAGTTGGTGGTATTTGACAC |
| sgPrkag1#2 | CACCGAGACTTAGACATGAATTC | AAACGAATTCATGTCTAAGTCTC |
| sgCops5#1 | CACCgTGTGTACTAACATCAATCC | AAACGGATTGATGTTAGTACACAc |
| sgCops5#2 | CACCgAATCCTGGCGGCGAAACCC | AAACGGGTTTCGCCGCCAGGATTc |
| sgPsmf1#1 | CACCgATCCTTAGACTCATACCGG | AAACCCGGTATGAGTCTAAGGATc |
| sgPsmf1#2 | CACCgCAAACGGCTACTATGCCTT | AAACAAGGCATAGTAGCCGTTTGc |
| sgPsma7#1 | CACCGTATCTCGGCCCTAATTGT | AAACACAATTAGGGCCGAGATAC |
| sgPsma7#2 | CACCgATACAGACGTTATCGTCCA | AAACTGGACGATAACGTCTGTATc |
| sgEwsr1#1 | CACCgATGGACAACAGAGTAGCTA | AAACTAGCTACTCTGTTGTCCATc |
| sgEwsr1#2 | CACCgCTACCATCAAACCATTCCA | AAACTGGAATGGTTTGATGGTAGc |
| sgPrkcd#1 | CACCgAGATTATCGGCCGCTGCAC | AAACGTGCAGCGGCCGATAATCTc |
| sgPrkcd#2 | CACCGAGTCTGTCGGAATATACC | AAACGGTATATTCCGACAGACTC |
| sgMapkap1#1 | CACCGCAGTACACGAGTGAAGGAC | AAACGTCCTTCACTCGTGTACTGC |
| sgMapkap1#2 | CACCgCTTGGAGTACTCATTAAAG | AAACCTTTAATGAGTACTCCAAGc |
| sgErcc2#1 | CACCGCAGACTCGGGCCAGCGTCA | AAACTGACGCTGGCCCGAGTCTGC |
| sgErcc2#2 | CACCgCGCTGATAGGCCACAATCA | AAACTGATTGTGGCCTATCAGCGc |
| sgHdac2#1 | CACCgCTTGATATACTCATCGCTG | AAACCAGCGATGAGTATATCAAGc |
| sgHdac2#2 | CACCgCATCTATACCATCTCTCAT | AAACATGAGAGATGGTATAGATGc |
| sgSetd2#1 | CACCgCTGCATTCGCTTAATATCC | AAACGGATATTAAGCGAATGCAGc |
| sgSetd2#2 | CACCGTATGTATCTTCCCGATGT | AAACACATCGGGAAGATACATAC |
| sgBrd1#1 | CACCGGGATATACGGAGCGCACA | AAACTGTGCGCTCCGTATATCCC |
| sgBrd1#2 | CACCGGCTCAATGAATACCGTGT | AAACACACGGTATTCATTGAGCC |
| sgTpp2#1 | CACCGATACACGGCTAAGCACTA | AAACTAGTGCTTAGCCGTGTATC |
| sgTpp2#2 | CACCGTTACTGTTGGAAATAACCG | AAACCGGTTATTTCCAACAGTAAC |
| sgUsp10#1 | CACCgTCGTGAGAGATATCCGCCC | AAACGGGCGGATATCTCTCACGAc |
| sgUsp10#2 | CACCgAGTCATCGAACCTAGTGAG | AAACCTCACTAGGTTCGATGACTc |
| sgRpa3#1 | CACCGTTACCACAGTATATCGAC | AAACGTCGATATACTGTGGTAAC |
| sgRpa3#2 | CACCGCTTGACGAGGAAATCTCT | AAACAGAGATTTCCTCGTCAAGC |
| sgPbk#1 | CACCGGCTCAGTACCAATATAAC | AAACGTTATATTGGTACTGAGCC |
| sgPbk#2 | CACCGAATGACTTAATAGAAGAG | AAACCTCTTCTATTAAGTCATTC |

| | | |
|---|---|---|
| sgCsnk2a1#1 | CACCGGACATGACAATTATGATC | AAACGATCATAATTGTCATGTCC |
| sgCsnk2a1#2 | CACCGACATTGTAAAAGACCCTG | AAACCAGGGTCTTTTACAATGTC |
| sgTaf7#1 | CACCgAGAATATGCCGCTACGGTG | AAACCACCGTAGCGGCATATTCTc |
| sgTaf7#2 | CACCGAAGCTGTCAGTACTCGTT | AAACAACGAGTACTGACAGCTTC |
| sgSmc4#1 | CACCgAGCATGGAATCAATAACAT | AAACATGTTATTGATTCCATGCTc |
| sgSmc4#2 | CACCgAGAAGCAGTAGTTAAGTTA | AAACTAACTTAACTACTGCTTCTc |
| sgPttg1ip#1 | CACCgCAGCATACGCCCCAGCGAG | AAACCTCGCTGGGGCGTATGCTGc |
| sgPttg1ip#2 | CACCCGCAGGAACCTCCGAGAGT | AAACACTCTCGGAGGTTCCTGCG |
| sgFancb#1 | CACCGTCACCGATCCACTCAATAG | AAACCTATTGAGTGGATCGGTGAC |
| sgFancb#3 | CACCGCAGCTGGTCTCTAGTAGCG | AAACCGCTACTAGAGACCAGCTGC |
| sgFanci#3 | CACCGAAATGGTTAAGTTAGACC | AAACGGTCTAACTTAACCATTTC |
| sgBrip1#2 | CACCGCAGACGAACTCTATCACG | AAACCGTGATAGAGTTCGTCTGC |
| sgPus7#1 | CACCGTACGTCGTAAAGCATGAGC | AAACGCTCATGCTTTACGACGTAC |
| sgPus7#3 | CACCGTAGTGCATAAACTCGCCGC | AAACGCGGCGAGTTTATGCACTAC |
| sgUbe2c#1 | CACCGTATAGCACACATCTCGTAG | AAACCTACGAGATGTGTGCTATAC |
| sgUbe2c#3 | CACCGCTGTTAAGAAAGATCACGG | AAACCCGTGATCTTTCTTAACAGC |
| sgUbe2t#3 | CACCGTCCAGGTTTGTGTCGCTGT | AAACACAGCGACACAAACCTGGAC |
| sgCtr | CACCGAGTTCACCGGCGTCATCGTC | AAACGACGATGACGCCGGTGAACTC |
| sgTRIP13#1 | CACCGTTGTCATCACATAATCG | AAACCGATTATGTGATGACAAC |
| sgTRIP13#2 | CACCGTGCATGCACTCAAATCGAT | AAACATCGATTTGAGTGCATGCAC |
| sgTRIP13#3 | CACCGGGCTAACGCTTTACACA | AAACTGTGTAAAGCGTTAGCCC |

**Supplementary Table 3.** GSEA results of global gene expression profiling after modulation of fetal and adult NUP98::KDM5A (related to Supplementary Fig. 1c)

| **NUP98::KDM5A vs EV fetal** | | | | |
|---|---|---|---|---|
| # | Gene set | NES | NOM p-val | FDR q-val |
| 1 | HOXA_cluster | 2.04 | 0.000 | 0.000 |
| 2 | HOXB_cluster | 1.33 | 0.016 | 0.080 |
| 3 | HOXA9_ZHANG | 2.22 | 0.000 | 0.000 |
| 4 | KLUSMANN_ARRAYSTAR_HSC_CB VS ALL_TOP200 | 1.81 | 0.000 | 0.007 |
| 5 | KLUSMANN_ARRAYSTAR_MONO | -2.52 | 0.000 | 0.000 |
| 6 | MARKEY_RB1_CHRONIC_LOF_DN | -2.11 | 0.000 | 0.000 |
| **NUP98::KDM5A vs EV Adult** | | | | |
| # | Gene set | NES | NOM p-val | FDR q-val |

| | | | | |
|---|---|---|---|---|
| 1 | HOXA_cluster | 2.600 | 0.000 | 0.000 |
| 2 | HOXB_cluster | 2.100 | 0.001 | 0.001 |
| 3 | HOXA9_ZHANG | 2.050 | 0.000 | 0.002 |
| 4 | KLUSMANN_ARRAYSTAR_HSC_CB VS ALL_TOP200 | 1.530 | 0.018 | 0.051 |
| 5 | KLUSMANN_ARRAYSTAR_MONO | -2.250 | 0.000 | 0.000 |
| 6 | MARKEY_RB1_CHRONIC_LOF_DN | -1.640 | 0.007 | 0.119 |

**Supplementary Table 4.** GSEA results of global gene expression profiling of fetal and adult NUP98::KDM5A (related to Fig. 1f)

| # | Gene set | NES | NOM p-val | FDR q-val |
|---|---|---|---|---|
| 1 | LTST FL VS LTST BM UP | 2.60 | 0 | 0 |
| 2 | FL VS BM UP | 2.40 | 0 | 0 |
| 3 | LTST FL VS LTST BM DOWN | -3.06 | 0 | 0 |
| 4 | FL VS BM DOWN | -3.07 | 0 | 0 |

**Supplementary Table 5.** GSEA results of global gene expression profiling after modulation of Trip13 in fetal NUP98::KDM5A (related to Fig. 3e)

| # | Gene set | NES | NOM p-val | FDR q-val |
|---|---|---|---|---|
| 1 | HALLMARK_P53_PATHWAY | 2.09 | 0.000 | 0.000 |
| 2 | REACTOME_STABILIZATION_OF_P53 | 2.03 | 0.000 | 0.000 |
| 3 | REACTOME_TP53_REGULATES_METABOLIC_GENES | 1.72 | 0.002 | 0.031 |
| 4 | KEGG_P53_SIGNALING_PATHWAY | 1.65 | 0.007 | 0.041 |
| 5 | REACTOME_TP53_REGULATES_TRANSCRIPTION_OF_CELL_DEATH_GENES | 1.62 | 0.012 | 0.046 |

**Supplementary Table 6.** GSEA results of global gene expression profiling after pharmacological modulation of Trip13 using DCZ0415 in fetal NUP98::KDM5A (related to Supplementary Fig. 5d)

| # | Gene set | NES | NOM p-val | FDR q-val |
|---|---|---|---|---|
| 1 | GOBP_DNA_DAMAGE_RESPONSE_SIGNAL_TRANSDUCTION_ BY_P53_CLASS_MEDIATOR_RESULTING_IN_CELL_CYCLE_ARREST | 1.81 | 0.002 | 0.027 |
| 2 | HALLMARK_P53_PATHWAY | 1.80 | 0.000 | 0.015 |
| 3 | WP_TP53_NETWORK | 1.58 | 0.036 | 0.056 |
| 4 | KEGG_P53_SIGNALING_PATHWAY | 1.55 | 0.021 | 0.050 |
| 5 | BIOCARTA_P53HYPOXIA_PATHWAY | 1.21 | 0.211 | 0.412 |
